# Conformation-dependent influences of hydrophobic amino acids in two in-register parallel *β*-sheet amyloids, an *α*-synuclein amyloid and a local structural model of PrP^Sc^

**DOI:** 10.1101/758938

**Authors:** Hiroki Otaki, Yuzuru Taguchi, Noriyuki Nishida

## Abstract

Prions are unconventional pathogens that encode the pathogenic information in conformations of the constituent abnormal isoform of prion protein (PrP^Sc^), independently of the nucleotide genome. Therefore, conformational diversity of PrP^Sc^ underlies the existence of many prion strains and species barriers of prions, although the conformational information is extremely limited. Interestingly, differences between polymorphic or species-specific residues responsible for the species/strain barriers are often caused by conservative replacements between hydrophobic amino acids. This implies that subtle differences among hydrophobic amino acids are significant for PrP^Sc^ structures. Here, we analyzed the influence of different hydrophobic residues on the structures of an in-register parallel *β*-sheet amyloid of *α*-synuclein (*α*Syn) using molecular dynamics (MD) simulation, and applied the knowledge from the *α*Syn amyloid to modeling a local structure of human PrP^Sc^ encompassing residues 107–143. We found that mutations equivalent to polymorphisms that cause transmission barriers substantially affect the stabilities; for example, the G127V mutation, which makes the host resistant to various human prion diseases, greatly destabilized the model amyloid. Our study demonstrates specifically how and in what structures hydrophobic residues can exert unique effects on in-register parallel *β*-sheet amyloids and provides insights into the molecular mechanism of the strain diversity of prions and other pathogenic amyloids.

**For Table of Contents Only:** 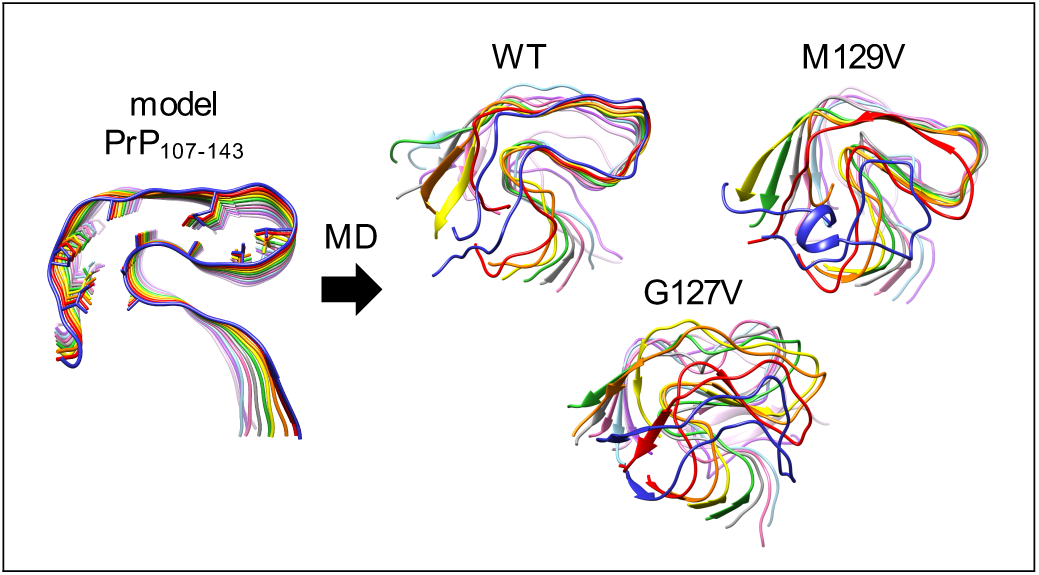

## 1 Introduction

Prion diseases are a group of neurodegenerative disorders that are characterized by the accumulation of the abnormal isoform (PrP^Sc^) of prion protein (PrP) in the central nervous system. ^1^ Prion diseases have three etiologies, sporadic, inherited, and acquired, depending on how the causative PrP^Sc^ propagation is initiated in the body. In sporadic and inherited prion diseases—for example, sporadic Creutzfeldt-Jakob disease (CJD) and fatal familial insomnia (FFI), respectively—the causative PrP^Sc^ are generated by the spontaneous conformational conversion of the endogenous normal isoform PrP (PrP^C^) into PrP^Sc^, with or without the aid of pathogenic mutations in the *PRNP* gene. Acquired prion diseases are caused by the intake of exogenous PrP^Sc^ as infectious prion agents, such as epidemic bovine spongiform encephalopathy (BSE) in cattle ^2^ and chronic wasting disease (CWD) in cervids.^3^ Prions behave similarly to viruses, with high infectivity, the existence of many strains, species/strain barriers, and adaptation to new hosts, despite the lack of conventional genetic material. These virus-like pathogenic properties of prions are hypothesized to be enciphered in the conformations of PrP^Sc^, a theory known as the protein-only hypothesis.^1, 4^

Indeed, the pathogenic properties of prions and the consequent clinical phenotypes are greatly affected by the primary structure of the constituent PrP^Sc^, which is consistent with the fundamental fact that the conformations of proteins are determined by their primary structures. For example, in sporadic CJD, PrP deposition patterns, lesion profiles in the brain, clinical presentations, and the apparent molecular sizes of protease-resistant cores of PrP^Sc^ vary depending on whether the polymorphic codon 129 is methionine (M129) or valine (V129).^5^ A pathogenic mutation D178N of human PrP causes either FFI or familial CJD in association with M129 or V129, respectively. ^6^ The vast majority of the new-variant CJD cases that resulted from the trans-species transmission of BSE to humans have been homozygous for M129.^7, 8^ Moreover, the methionine/leucine polymorphism at codon 132 of elk PrP, which is equivalent to the codon 129 polymorphism of human PrP, also affects the susceptibility to CWD. ^9, 10^ From the viewpoint of the protein-only hypothesis, polymorphisms could affect the stability of PrP^C^/PrP^Sc^, and should also affect disease susceptibility through structural alterations of the constituent PrP^Sc^; however, the specific manner how the subtle differences between those hydrophobic amino acids affect the structures of PrP^Sc^ has not been identified yet. Detailed structures of PrP^Sc^ are needed for such investigation, but even whether PrP^Sc^ is an in-register parallel *β*-sheet amyloid or a *β*-solenoid has been controversial due to its incompatibility with conventional high-resolution structural analyses. ^11–19^ In 2021, while our study was under revision, a high-resolution structure of a fully infectious, brain-derived prion was solved using cryo-electron microscopy (cryo-EM) by Kraus et al.^20^ This PrP^Sc^ (263K strain) has an in-register parallel *β*-sheet architecture without paired protofibrils. Then, cryo-EM structures of the glycosylphosphatidylinositol (GPI)-anchored and anchorless mouse-adapted RML scrapie strains proved that the conformations of PrP^Sc^ differ depending on the strains. ^21, 22^

In addition to the well-known example of methionine/valine polymorphic codon 129, strain/species barriers of prions can also be caused by conservative replacements between different hydrophobic amino acids at other residues. In experiments with a C-terminally-truncated Y145Stop mutant of human PrP and the counterparts of mouse and Syrian hamster PrPs, whether residue 138 or 139 (in human numbering throughout, unless otherwise noted) was isoleucine or methionine was critical for efficient amyloid formation and cross-seeding among the PrPs. ^23, 24^ In the transmission of the prions of sporadic CJD homozygous for M129 to transgenic mice expressing human-mouse chimeric PrP, I138M substitution substantially extended the incubation periods. ^25^ Different hydrophobic amino acids at codons 109 and 112 influenced the transmission efficiencies of prions among different hamster species. ^26, 27^ In in vitro cross-seeding between ovine PrP and cervine CWD, an I208M (in ovine numbering) mutation showed a profound influence on the seeding efficiencies. ^28^

Given these documented facts, we reasoned that the subtle differences of hydrophobic side chains are influential because generally amyloids need hydrophobic cores for stabilization; thus we consider studying the effects of the replacement of hydrophobic amino acids with different ones on the structures of amyloids is important, particulary whether methionine has unique properties compared with other hydrophobic amino acids. Currently available structures of pathogenic in-register parallel *β*-sheet amyloids share basic structures; they consist of intramolecular pairs of short *β*-strands and at least one *β*-arch. We reasoned that pairing of the constituent *β*-strands may be required for stability of an in-register parallel *β*-sheet amyloid and, in the case, stable *β*-arches which bend the backbone at 180 degrees would be advantageous or even essential for efficient intramolecular coupling of the *β*-strands, and eventually efficient conversion into the amyloid form of the molecule. For instance, it is conceivable that a peptide molecule with a (or more) region prone to stable 180-degree-bending *β*-arch in the middle of the molecule may be more susceptible to conformational conversion into in-register parallel *β*-sheet amyloid, because of efficient intramolecular pairing of *β*-strands. Therefore, we are interested in determinants of stability of 180-degree-bending *β*-arches, particularly U-shaped *β*-arch. The present study is aimed to identify some of the conditions which stabilize or destabilize *β*-arches.

We used an in-register parallel *β*-sheet amyloid of *α*-synuclein (*α*Syn) as a surrogate local structural model for PrP^Sc^, as in our previous studies.^29, 30^ *α*Syn amyloid is the main component of Lewy bodies, which is a hallmark of Parkinson’s disease and dementia with Lewy bodies (DLB), and reportedly has prion-like properties which include transmissibility and strain diversity. ^31^ For instance, *α*Syn forms various types of amyloids in vitro, which are different in appearance, proteolytic fragment patterns, and cytotoxicity.^32^ Moreover, *α*Syn amyloids isolated from patients with DLB and multiple system atrophy had different proteolytic fragment patterns suggestive of distinct conformations. ^33^ These findings are highly reminiscent of prions and imply that prion-like properties are inherent in in-register parallel *β*-sheet structures. Detailed structures of *α*Syn amyloids have been determined using solid-state NMR (ssNMR)^34^ or cryo-EM;^35, 36^ thus in the present study we first used a Greek-key *α*Syn amyloid (PDB ID: 2N0A;^34^ Figure 1) to investigate the effects of different hydrophobic residues on the in-register parallel *β*-sheet structures. ^37^ These experiments revealed that the lengths and C*_β_*-branching of the side chains of hydrophobic residues are important for the stability of local amyloid structures, particularly in a U-shaped *β*-arch. We then applied the knowledge from the *α*Syn amyloid to a local structural model of PrP^Sc^ encompassing residues 107 to 143 of human PrP, PrP_107-143_, under the assumption that PrP^Sc^ was also an in-register parallel *β*-sheet amyloid. Specifically, we assessed how mutations and polymorphisms associated with the strain diversity of prions affected the structures of the model amyloid using molecular dynamics (MD) simulation. The results of these studies demonstrated how different types of hydrophobic amino acids specifically affect the structures of in-register parallel *β*-sheet amyloids of *α*Syn and PrP_107-143_. They also showed that the structural stability of U-shaped *β*-arches requires hydrophobic cores with well-balanced interactions among the constituent hydrophobic residues.

**Figure 1:**
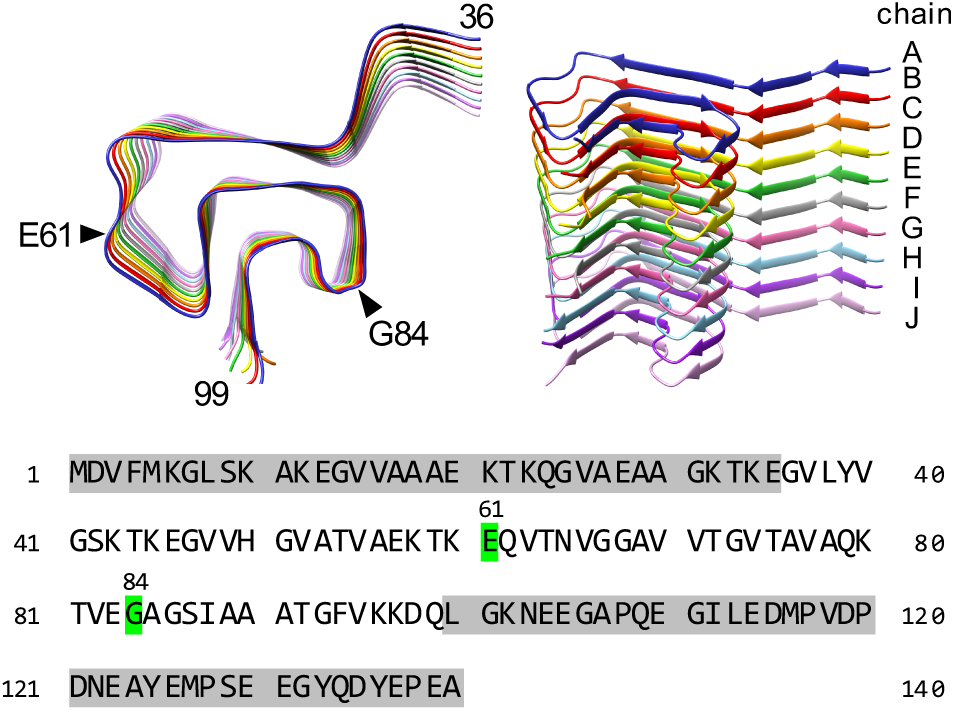
(Top) Structure of the Greek-key *α*Syn amyloid (PDB ID: 2N0A), in which the N- and C-terminal-side regions (residues 1–35 and 100–140, respectively) are truncated. The black triangles indicate the positions of residues E61 and G84. The chains of the amyloid stack are referred to as chains A–J. (Bottom) Amino acid sequence of *α*Syn. The truncated residues are highlighted in gray. The residues E61 and G84 are highlighted in green.

## 2 Results and Discussion

### 2.1 Investigation of the mechanism by which G84M mutation stabilizes the ***α***Syn amyloid

In a previous study, we compared the effects of isoleucine substitutions at residues 61 (E61I) in a flat-loop (loop(57–62)) and 84 (G84I) in a U-shaped loop (loop(84–87)) in the *α*Syn amyloids.^29^ We showed that the G84I mutant substantially destabilized the U-shaped loop, whereas the E61I mutant had little influence on the stability of the *α*Syn amyloids. Here, we investigated the influences of hydrophobic amino acids on the local structures of *α*Syn amyloids by introducing methionine substitutions at the same positions (E61M and G84M). For all the model fibrils, the radius of gyration fluctuated around 24.5 Å during the simulation (Figure S1), which indicated that the simulation time was too short to discuss (de)stabilization of the whole structure of the *α*Syn fibrils. However, our simulation highlighted the (de)stabilization in the local structure of the fibrils as shown below.

Figure 2A shows the final snapshots of MD simulations of *α*Syn amyloids, and Figure 2B shows the heatmaps of the *β*-sheet propensity. The MD simulations of homo-oligomer amyloids of *α*Syn(E61M) and *α*Syn(G84M) revealed that both the mutations tended to stabilize the local structures, particularly when these amyloids induced new *β*-sheets in the flat-loop for the E61M mutant, or U-shaped loop for the G84M mutant (Figure 2A). This stabilization in *α*Syn(E61M) could be interpreted as an effect of the charge difference in the E61M mutation: the side chains of M61 formed hydrophobic contacts with those of the neighboring chains along the stacking direction in the E61M mutant, whereas the negative charge of glutamate hampered the stabilization between the neighboring chains in the wild-type (WT). Overall, the influence of the methionine substitution (E61M) was similar to that of the isoleucine substitution (E61I) in the flat-loop.

**Figure 2:**
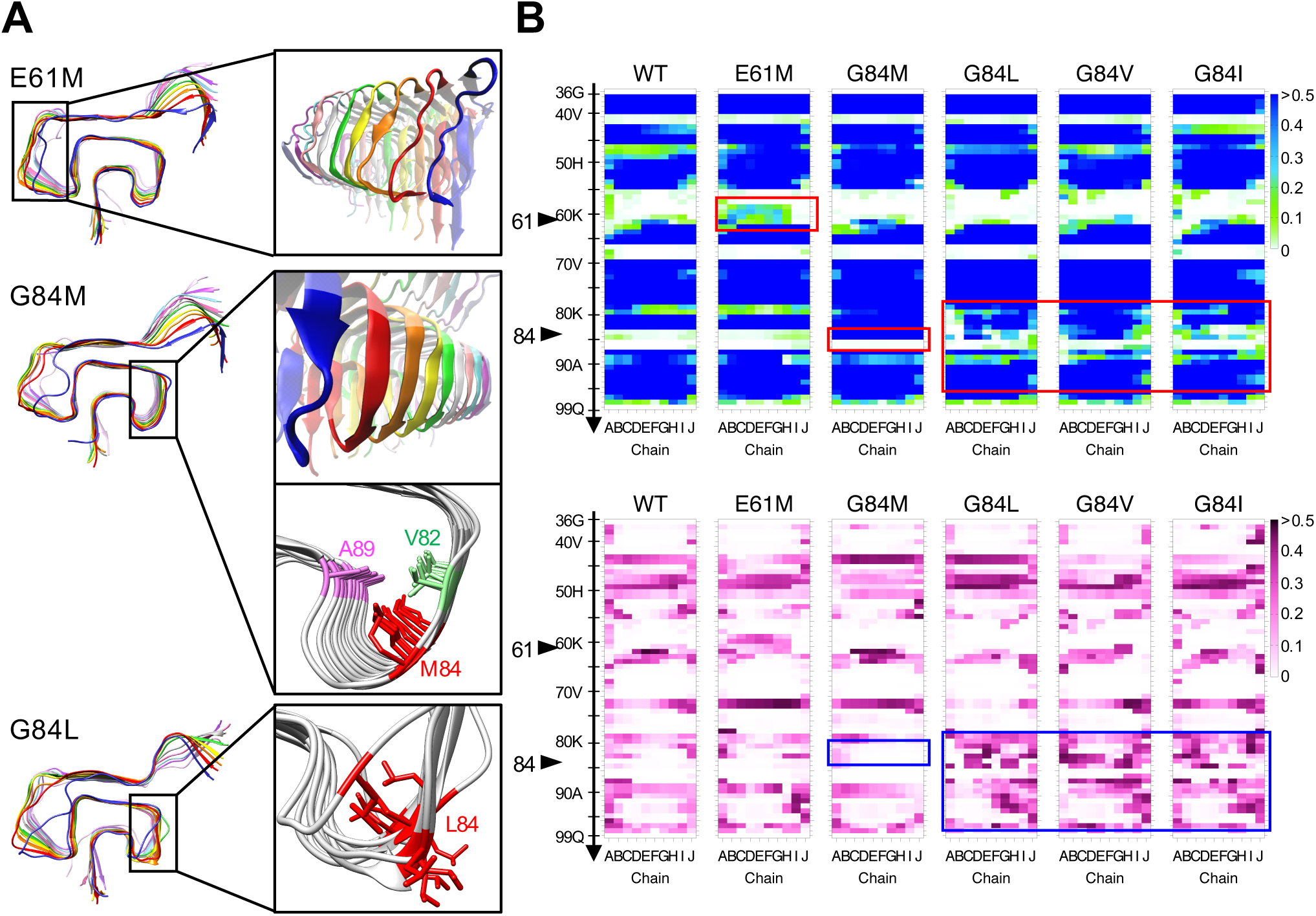
Influences of E61M and G84M/L/V/I substitutions on the local structures of Greek-key *α*Syn amyloid: (A) Final snapshots of *α*Syn(E61M), *α*Syn(G84M), and *α*Syn(G84L) amyloids after 400-ns MD simulations. The insets present magnified views of the indicated regions. Note that the *β*-sheets are induced in loop(57–62) of the *α*Syn(E61M) amyloid and that the *α*Syn(G84M) amyloid has *β*-sheets extended to loop(84–87). In *α*Syn(G84M) and *α*Syn(G84L), the directions of the side chains of residue 84 after a 400-ns MD simulation are shown with red sticks in the insets. In the inset of *α*Syn(G84M), the side chains of V82 and A89 are also shown with green and pink sticks, respectively. They are close to the side chains of M84 and form hydrophobic contacts (see also Figure 3). In *α*Syn(G84L), the local structures around the mutation are disturbed. Some side chains of L84 are pointing outward, whereas those of M84 remain inside of *α*Syn(G84M). (B) Heatmaps of the average *β*-sheet propensity values (Avg-*β*) and the standard deviation of the values (SD-*β*) based on five (for WT) and three (for the mutants) independent 400-ns MD simulations. The vertical axes and horizontal axes of all the heatmaps represent the residue numbers and the chains A–J, respectively. The black triangles indicate the positions of residues 61 and 84. Avg-*β* values in loop(57–62) of *α*Syn(E61M) and loop(84–87) of the *α*Syn(G84M) amyloids increased (E61M and G84M, red boxes), corresponding to the induced *β*-sheets in those regions (see (A)). The relatively low SD-*β* values in the *α*Syn(G84M) amyloid (G84M, blue box) reflect the stabilized local structures. In *α*Syn(G84L), *α*Syn(G84V), and *α*Syn(G84I), the structural disturbances caused by the mutation are reflected to the heatmap (G84L/V/I, red and blue boxes). The heatmaps of the WT and G84I mutant are reprinted with permission under the CC BY 4.0 license from Taguchi et al.^29^ Copyright 2019 MDPI.

The effects of the G84M substitution were more impressive: almost the entire region C-terminal to residue 80, including loop(84–87), was converted to stable *β*-sheets, unlike in the case of the G84I substitution. ^29^ We thus further compared the substitution effects of G84M, G84L, G84V, and G84I. In the G84L mutant (*α*Syn(G84L)), the loop region around L84 was substantially destabilized like that of *α*Syn(G84I).^29^ A G84V mutant (*α*Syn(G84V)) also showed a *β*-sheet propensity similar to that of *α*Syn(G84L). Because we were interested in the opposite effects of G84M and G84L/V/I, we focused on residue 84 to identify the underlying mechanisms.

The directions of the side chains of *α*Syn(G84M) and *α*Syn(G84L) are compared in the insets of Figure 2A. The final status of 400-ns simulations was different between the *α*Syn(G84M) and *α*Syn(G84L) amyloids. All the side chains of methionine at residue 84 (M84) pointed inward of loop(84–87) and were close to those of V82 and A89, whereas some side chains of leucine at residue 84 (L84) flipped and pointed outward.

The hydrophobic contact diagrams are shown in Figure 3. The diagrams of the *α*Syn(G84M), *α*Syn(G84L), *α*Syn(G84V), and *α*Syn(G84I) amyloids clearly exhibits differences. M84 of the *α*Syn(G84M) amyloid had strong intra-chain interactions with residues 82 and 89 in all the layers, and the interactions were reproducible. In contrast, the *α*Syn(G84L) and *α*Syn(G84V) amyloids only infrequently showed that pattern or a pattern similar to the amyloids of another destabilizing mutation, G84I.^29^ This result was reflected in the distance between residues 84 and 89. The C*_α_*-C*_α_*distances (*d*C*_α_*) between the residues 84 and 89 were about 8.3 Å in all the chains during the simulations of *α*Syn(G84M), whereas the distances varied among chains and fluctuated during the simulations of *α*Syn(G84L), *α*Syn(G84V), and *α*Syn(G84I) (Figure S2).

**Figure 3:**
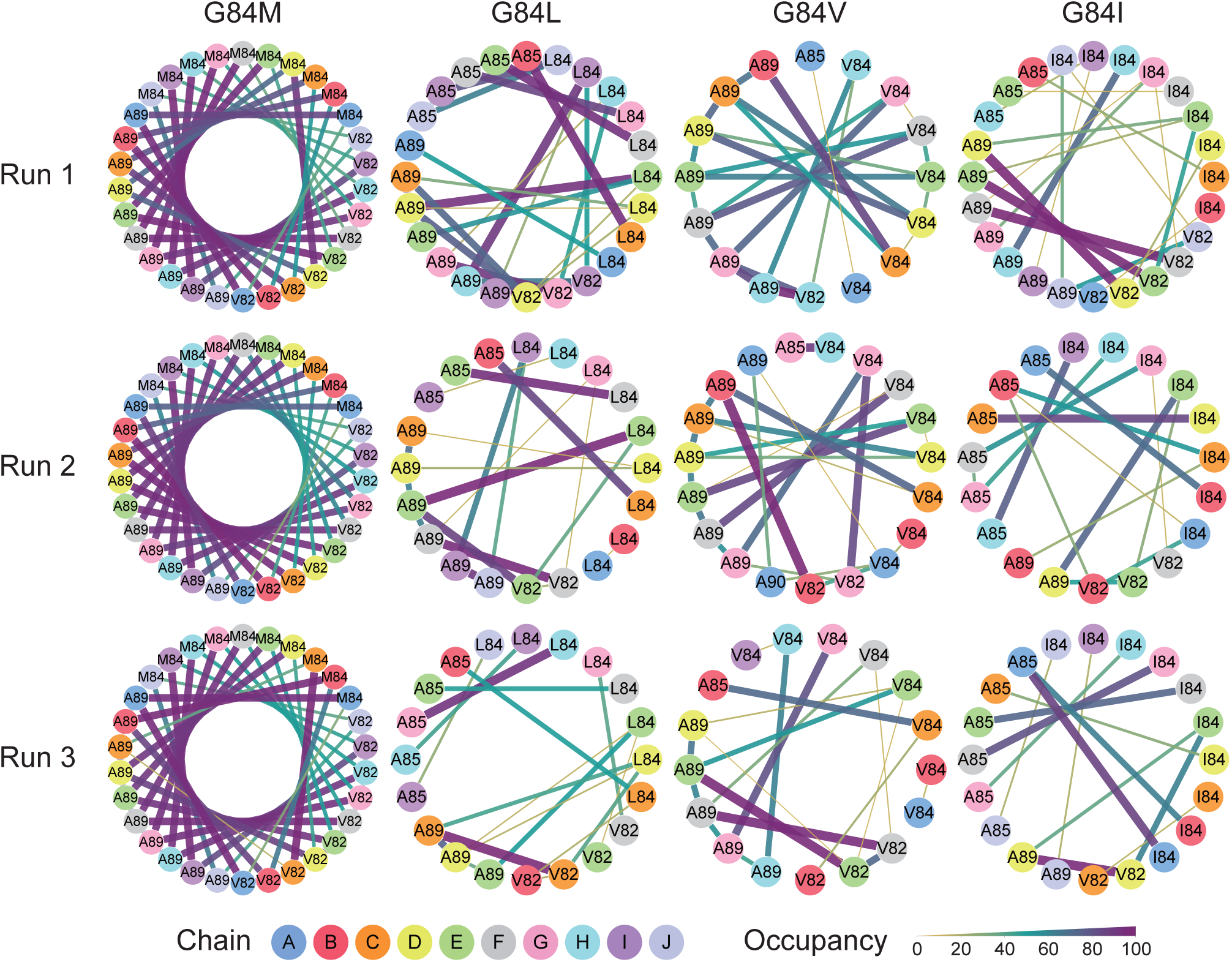
Hydrophobic contact diagrams based on 400-ns simulations of *α*Syn amyloids. Only the residues involved in the interactions with residue 84 are presented. Residue numbers and chains are indicated by the numbers and colors of the dots, respectively. The thickness and color of each line indicate the occupancy of the hydrophobic contact.

The unique stabilizing effects of M84 would be attributable to the long side chain, which can extend across the *β*-arch like a crossbeam. During the simulation, the ends of the side chains were fixed in the proximity of A89 (Movie S1). The intimate interactions with A89 reinforced the hydrophobic core and stabilized the *β*-arch. Leucine could not replace methionine despite their similar hydrophobicities and non-branching C*_β_* atoms, because its side chain was not long enough for stable interactions with A89. Indeed, other groups have confirmed that the subtle difference between methionine and leucine is significant in at least one case of prion propagation—namely, the methionine/leucine polymorphism of elk PrP affects the susceptibility to CWD infection.^9, 10^ The local structures of CWD-PrP^Sc^ around the polymorphic residue might be similar to the U-shaped loop of *α*Syn—that is, the long side chain of methionine may be essential for stability.

### 2.2 Modeling a local structure of PrP^Sc^ based on knowledge from the ***α***Syn amyloids

As in our previous studies on *α*Syn amyloids,^29, 30^ we hypothesized that PrP^Sc^ has an in-register parallel *β*-sheet structure and attempted to model a local structure of PrP^Sc^ with a U-shaped loop. (Note that the present study was conducted before the publication of the paper by Kraus et al. ^20^) A region comprising a glycine-rich motif encompassing residues 123–127 of human PrP (*−*GGL_125_GG*−*) seemed to be suitable for a U-shaped loop. In addition, the anti-prion effect of the valine at residue 127 against various sporadic CJD^38^ was reminiscent of the destabilizing effects of V84 in the *α*Syn amyloid, which suggested the presence of a U-shaped loop in the region. Moreover, the structures of amyloid cores of Y145Stop-mutant PrP investigated with ssNMR were available, although not in atomic resolution.^16, 39^ Amyloids of the Y145Stop-mutant induced infectious PrP^Sc^, which caused bona fide prion disease when inoculated into mice,^40^ and could therefore share similar local structures with full-length PrP^Sc^. We thus tentatively modeled the local structures of human CJD PrP^Sc^ in region 107–143, PrP_107-143_, utilizing the structural model of the Y145Stop mutant propounded by Theint et al. ^39^ and the knowledge from the *α*Syn amyloid for stable *β*-arches discussed in Section 2.1. That is, the side chains of hydrophobic residues in the U-shaped loop (A120, V122, L125, and M129) were modeled to point inward for investigation of their hydrophobic contact. The structure of the modeled PrP_107-143_ amyloid is represented in Figure 4A. The detailed procedure used for the modeling is described in Section 4.4. Although a *β*-solenoid structural model based on cryo-EM was recently reported, ^41^ the particular amyloid was not demonstrated to cause prion disease in mice and we did not adopt it. A comparison of our model and the cryo-EM structure ^20^ is given in Section 2.8. Recently, detailed structures of RML and 263K prions were reported. Although their structures in the relevant region are different from our model, they are mouse and hamster prion respectively. Furthermore, as the main purpose of our study is to investigate factors that contribute to stability of U-shaped *β*-arches, the discrepancy does not necessarily undermine the value of our research.

**Figure 4:**
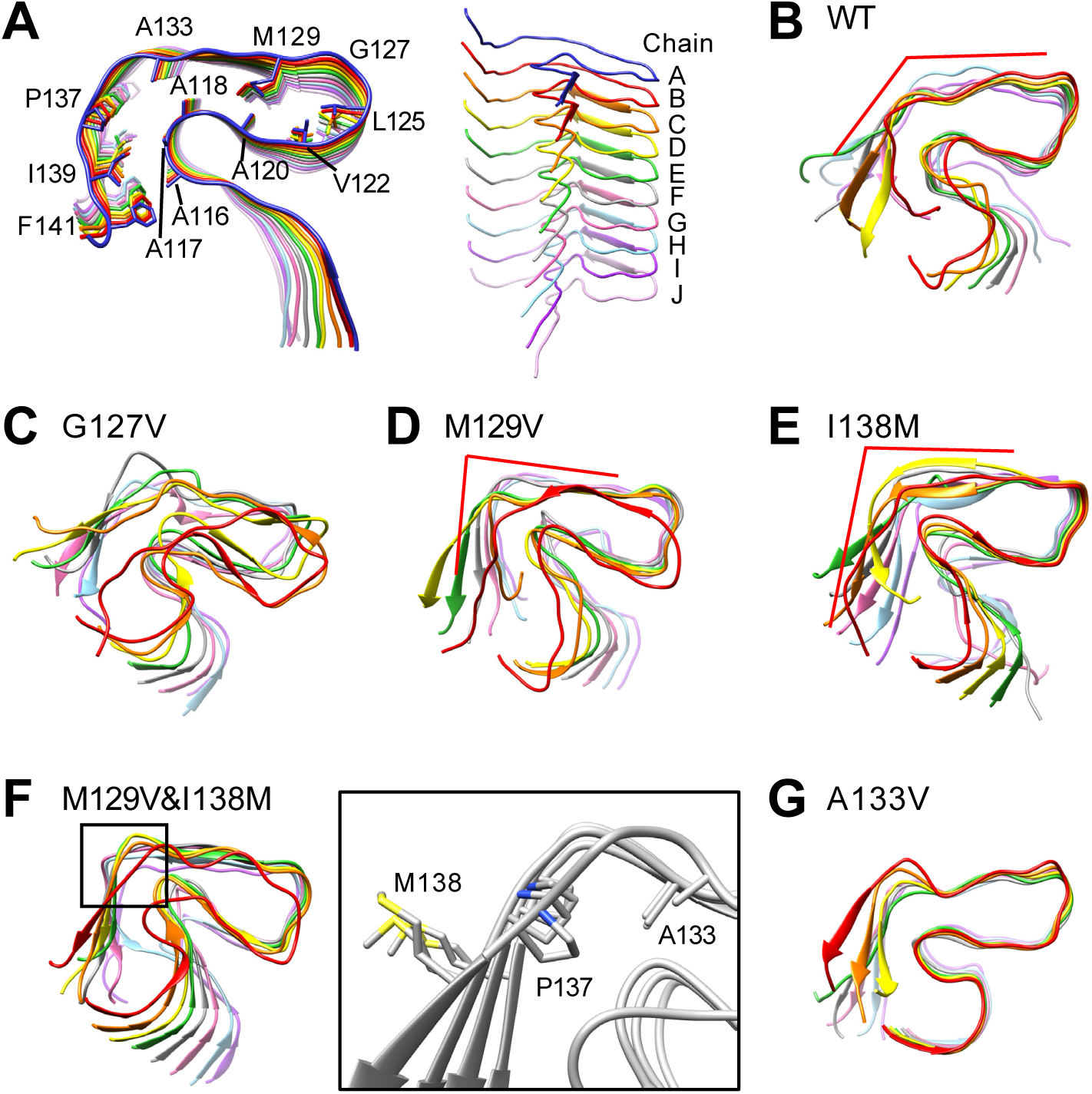
Local-structural model of PrP^Sc^, PrP_107-143_, and final snapshots of 400-ns MD simulations (Run 3). For all the snapshots of five runs, see Figure S3: (A) A PrP_107-143_ amyloid as a local-structural model of PrP^Sc^ and an oblique view showing the 10 layers (chains A–J). Side chains of the residues that are mentioned later are annotated. (B) A final snapshot of WT PrP_107-143_. (C) A final snapshot of PrP_107-143_(G127V). (D) A final snapshot of PrP_107-143_(M129V). (E) A final snapshot of PrP_107-143_(I138M). (F) (Left) A final snapshot of PrP_107-143_(M129V&I138M). The box indicates Ω-shaped loops encompassing A133 to P137. (Right) Positional relationships of the side chains of A133, P137, and M138 in chains B–E. (G) A final snapshot of PrP_107-143_(A133V). For (B)–(G), chains A and J are removed for clarity.

To sample the conformational space of PrP_107-143_ more broadly, we conducted five independent 400-ns MD simulations for the WT. Although the stack ends of the WT model amyloid (chains A and J) were not very stable as expected, as a whole our model was sufficiently stable during the MD simulations (Figures 4B, S3A, and S4A for the last snapshots; Figure S5 for the radius of gyration; see also Movie S2). We then modeled mutants of PrP_107-143_ (G127V, M129V, I138M, M129V&I138M, and A133V) and performed MD simulations. The results are discussed in the following sections.

### 2.3 Influence of G127V mutation on the PrP_107-143_ amyloid

A final snapshot of MD simulation for the G127V mutant, PrP_107-143_(G127V), is presented in Figure 4C, from which we confirmed the destabilizing effects of the mutation on the region including the U-shaped loop (residues 120–134; see also Figures S3B and S4B, and Movie S3). These effects are reflected in the root-mean-square fluctuation (RMSF) of C*_α_*atoms and *β*-sheet propensity values shown in Figure 5: the heatmaps of PrP_107-143_(G127V) indicate the disorder around the U-shaped loop. To validate the (in)stability of the modeled amyloids, we used a parameter defined by the sum total of the persistence of the hydrophobic contacts between two hydrophobic residues over the eight chains, B–I (hereafter called the proximity score; see Figure S6A). This score counts hydrophobic contacts between the two residues of interest irrespective of intra- or inter-chain interactions and presumably indicates the total contribution of these residues to the stability of the amyloid. Unlike when using contact maps, we can set a threshold for the contact so that the score reflects the stability of the system. Here, the threshold for the hydrophobic contact was set at 5 Å,^42^ because the typical distance between neighboring *β*-strands within a *β*-sheet was 4.8 Å. The scores correlated with distances between the two corresponding residues in certain cases (Figure S6B) and reflected the stability of the hydrophobic cores. Figures 6 and 7 summarize the proximity scores in the U-shaped loop and the N-/C-terminal-side regions, respectively. The proximity scores for V122-V122, L125-L125, and L130-L130 of PrP_107-143_(G127V) decreased significantly compared with those of the WT. The score for V122-M129 also decreased, although not to a statistically significant degree. These results indicated that those residues were frequently wide apart, consistent with disordered U-shaped loops. In the other regions, the proximity scores for A115-V121 and A118-P137 decreased, although this result was also not statistically significant. The V127 polymorphism of human PrP is known to confer the protective effects against CJD transmission.^38^ Further research (e.g., a longer MD simulation) will be needed to ascertain the relationship between the protective effect and the destabilization of the amyloid structure caused by the G127V mutation.

**Figure 5:**
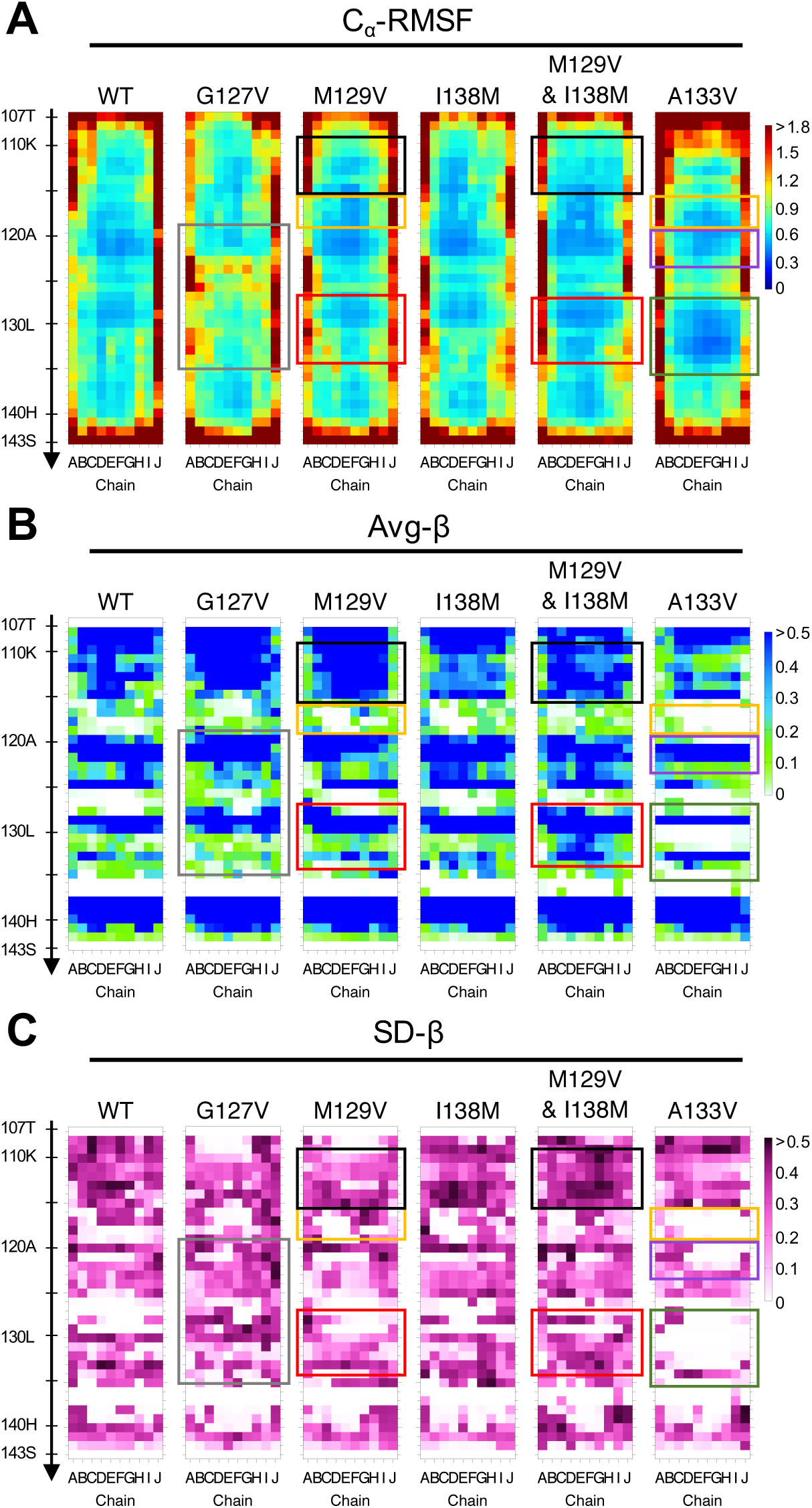
Heatmaps showing (A) the average of root-mean-square fluctuation (RMSF) of C*_α_* atoms, (B) the Avg-*β*, and (C) the SD-*β* of each residue of the WT and the mutant PrP_107-143_ amyloids. The values were calculated from five independent 400-ns MD runs: gray boxes, a disordered loop(120–134); yellow boxes, a more demarcated loop(116–119) than in the WT; black boxes, a more stable *β*-sheet(110–115) with higher *β*-sheet propensity and lower RMSF values than in the WT; red boxes, a more stable *β*-sheet(128–133) with lower RMSF and higher *β*-sheet propensity around L130 than in the WT. Higher SD-*β* values at A133 are induced by an “upward shift” (see Figure 8E,F); purple boxes, a more demarcated *β*-sheet(121–122) with lower RMSF values than in the WT; green boxes, a more demarcated *β*-sheet propensity at M129 and V133 with lower RMSF and SD-*β* values than in the WT.

**Figure 6:**
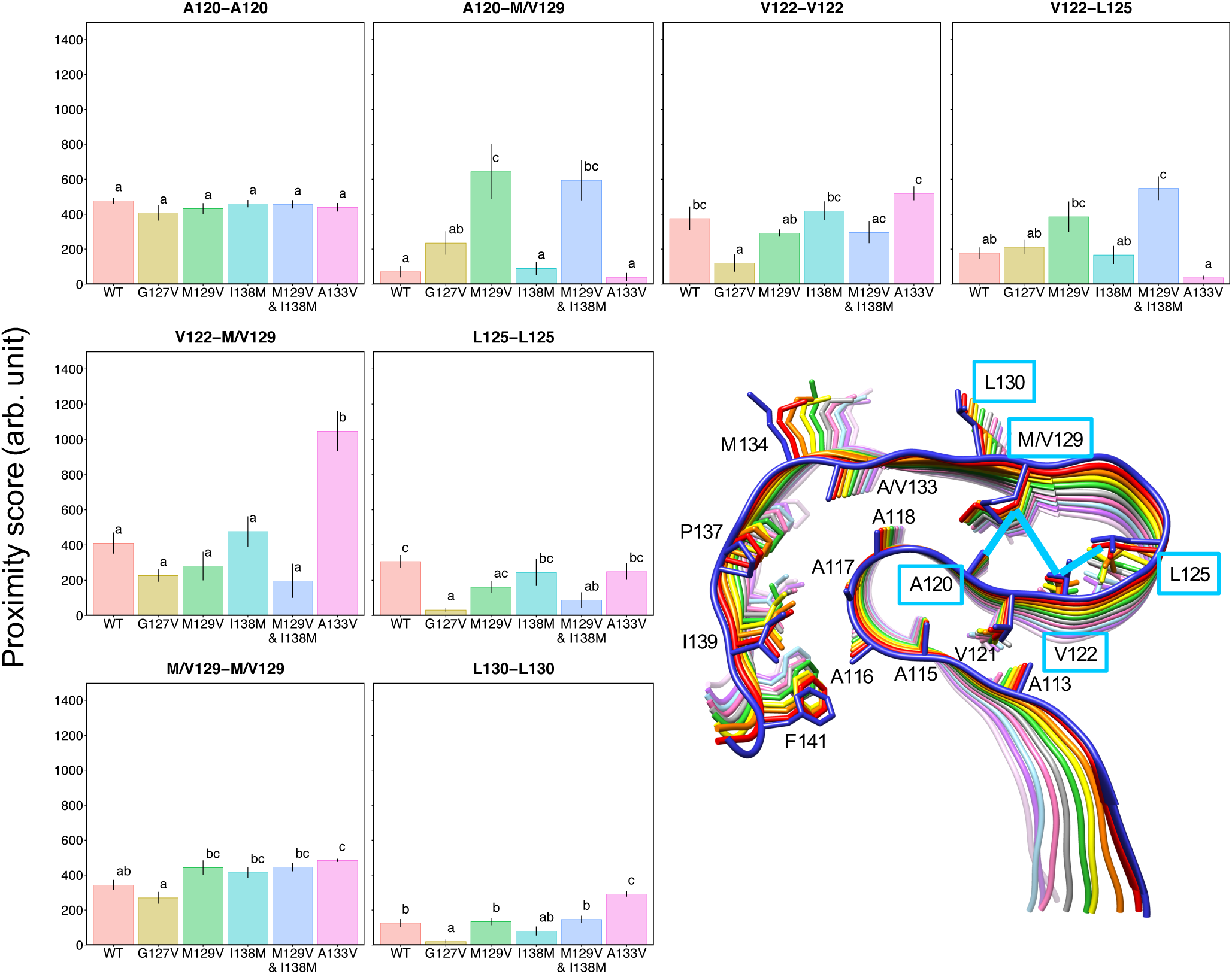
Bar plots of proximity scores between two residues in the U-shaped loop (residues 120–130). The proximity score was evaluated with chains B–I, because chains of the stack ends (i.e., chains A and J) were not stable during the MD simulations (see Figure S3). The error bar represents the standard error (SE) of the mean of values from five independent MD runs. Bars sharing the same letter are not significantly different according to Tukey’s test with *α* = 0.05. In the figure of the PrP_107-143_ amyloid, plotted interactions are represented with light-blue boxed residues (for hydrophobic contacts along the stacking direction) or lines (for the others).

**Figure 7:**
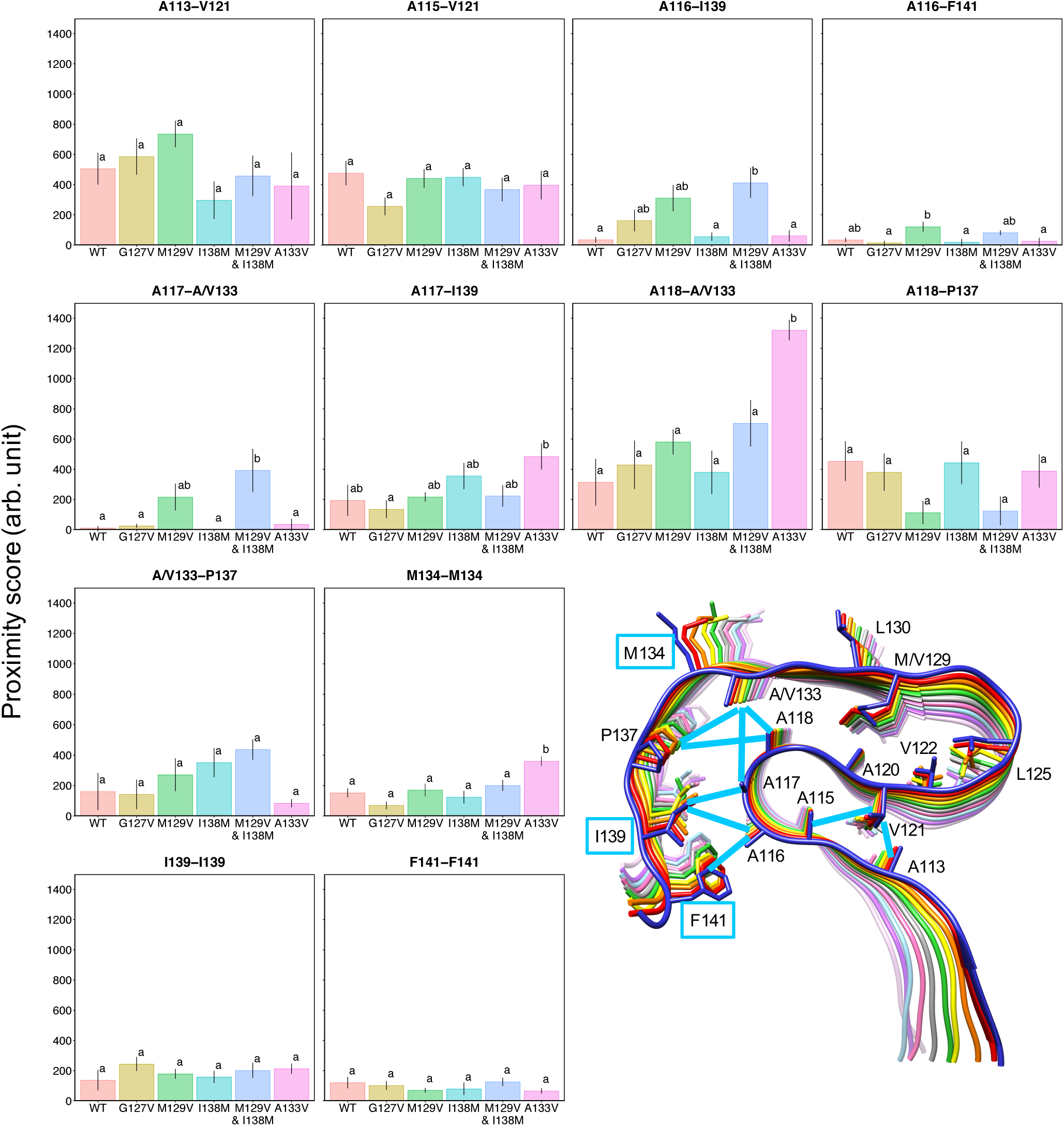
Identical to Figure 6, except for the residues in the N- and C-terminal-side regions.

### 2.4 Influence of M129V mutation on the PrP_107-143_ amyloid

Next, we analyzed the influence of M129V substitution, which is equivalent to V129 polymorphism of human PrP, on the stability of the hydrophobic cores. The mutant amyloid, PrP_107-143_(M129V), appeared more stable during the 400-ns simulation than the WT (Figure 4D). Comparison of the RMSF and *β*-sheet propensity values between the WT and M129V validated the stability, with more demarcated loops (residues 116–119, especially chains B–E) and more stable *β*-sheets (residues 110–115 and 128–133) (Figure 5). Notably, the C-terminal-side region tended to lean over toward the N-terminal-side region (Figures 4D, S3C, and S4C), presumably due to bending at the loop(134–137). Because of the bending, the angle between the *β*-sheet(128–133) and the *β*-sheet(139–141) appeared to be near-square to acute in many runs of the M129V mutant. The increase of the proximity score for A133-P137 supported this bending (Figure 7).

Proximity scores of the WT and the mutant revealed more detailed differences. Most notable was the alteration in the balance of interactions between A120, V122, L125, and V129 in the hydrophobic cores of the U-shaped loops (Figure 6). The WT showed outstandingly high intra-chain proximity scores for V122-M129 (*∼* 400) in the U-shaped loop, whereas the M129V mutation reduced the score (to about 250) and, instead, increased the scores for A120-V129 (to about 650) and V122-L125 (to about 400). The markedly enhanced hydrophobic interaction for A120-V129 may have yielded the higher *β*-sheet propensity of A120 (Figure 5B). The higher scores for A120-A120 and V129-V129 reflected the greater structural stability of the mutant amyloid. The mutation also affected interactions in remote regions, as demonstrated by the decrease of the proximity score for A118-P137 (Figure 7), although this decrease was not statistically significant. Some increases of the proximity scores were also seen, such as those for A113-V121, A116-I139, A116-F141, A117-A133, A118-A133, and A133-P137, although these increases were also not statistically significant. The altered interactions in the N-terminal-side region (A113-V121 and A115-V121) were particularly interesting, because they were located on the opposite side from V129 across *β*-strand 120– 122 and did not directly contact V129. Presumably, the dynamics of the *β*-sheet(120–122) were initially affected by the mutation, and subsequently altered the interaction patterns of V121; this mechanism could explain the increase in *β*-sheet propensity for residues 110–115. Notably, the above description almost applied to the M129V&I138M mutant, and some interactions were enhanced by the I138M mutation (see Section 2.5).

As shown in Figure 4D, the distance between *β*-sheet(120–122) and *β*-sheet(128–133) seemed to be narrower in the M129V mutant. Indeed, the M129V mutation increased the proximity scores for A118-A133 (Figure 7), and the distances between the C*_α_*atoms of A118 and S132, *d*C*_α_*(A118–S132), tended to be shorter in mutants with M129V mutations (Figure 8A, green and blue squares) than in the WT (red dots). Interestingly, the WT and mutants with M129 mostly had shorter *d*C*_α_*(A118–A/V133) than *d*C*_α_*(A118–S132), whereas *d*C*_α_*(A118–S132) was slightly shorter than *d*C*_α_*(A118–A/V133) in the V129 mutants (M129V and M129V&I138M) (Figure 8B). Figure 8C shows the correlation between *d*C*_α_*(A120–M/V129) and *d*C*_α_*(A118–S132). The mutants with V129 showed obviously shorter *d*C*_α_*(A120–M/V129) compared with the mutants with M129. Additionally, in the mutants with V129, the distances between C*_α_* atoms of A120 and V129, *d*C*_α_*(A120–V129), were shorter than *d*C*_α_*(V122–V129), while the other mutants showed the opposite tendency (Figure 8D). All these findings could be explained by assuming an “upward” positional shift of the *β*-sheet(128–133) in the mutants with V129 (Figure 8E,F) that is attributed to the side chain of V129 being shorter than that of M129. This shortness enabled V129 to further approach toward A120 and consequently caused well-balanced interactions of the hydrophobic residues and closer positioning of the two *β*-sheets. In addition to the local effects, the positional shift also expanded the range of motion of the C-terminal-side region, leading to a loss of interaction for A118-P137 and facilitating an interaction of A116-F141 (Figure 7). Notably, this “upward shift” was compatible with the bending at the loop(134–137) mentioned above. In contrast, in the case of the long side chain of M129, the biased interactions with V122 seemed to hamper the approach of residue 129 toward A120 (Figure 6, A120-M/V129 and V122-M/V129).

**Figure 8:**
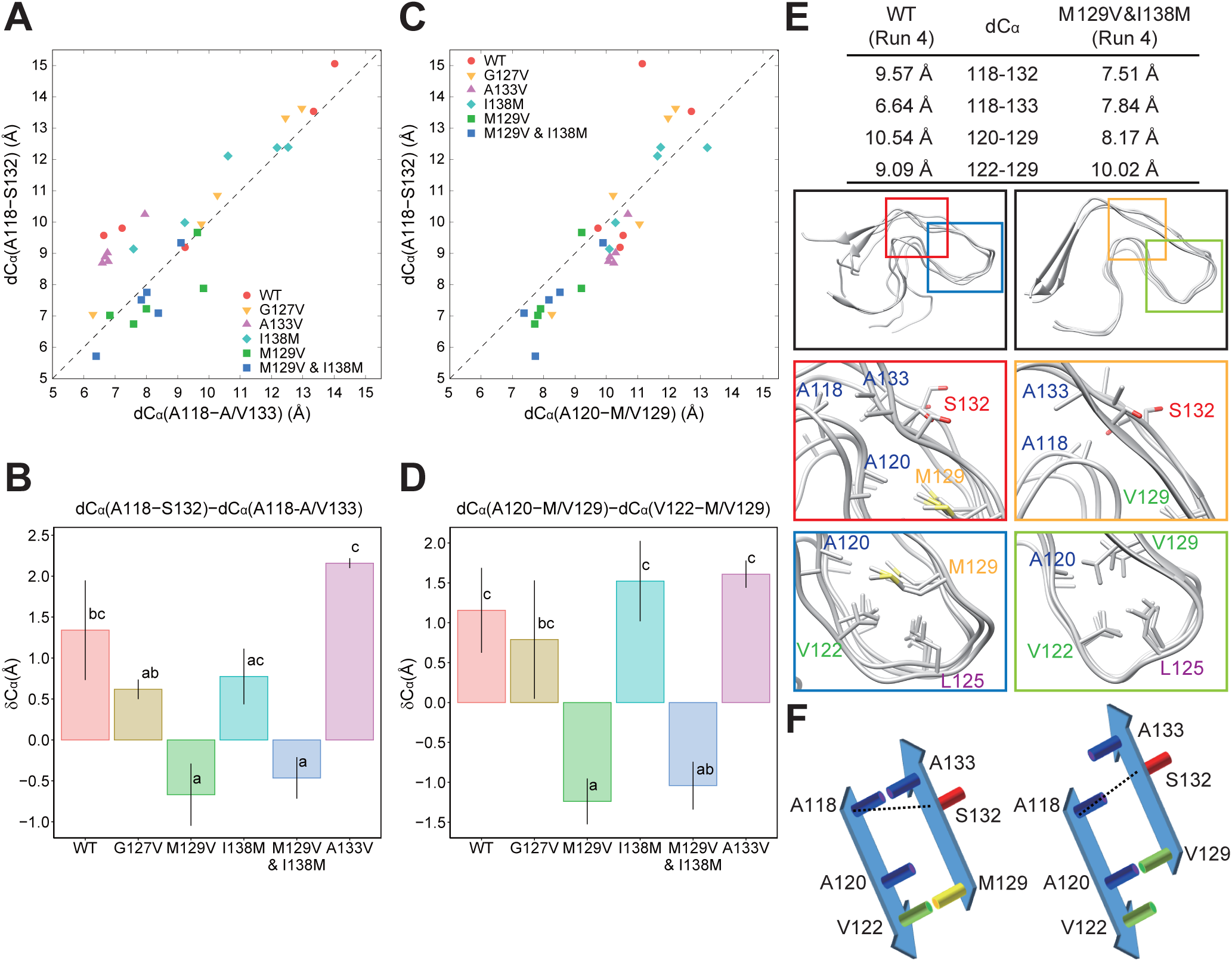
(A) Scatter plot showing correlations of the final C*_α_*-C*_α_*distances for A118-A/V133 (*d*C*_α_*(A118-A/V133)) and A118-S132 (*d*C*_α_*(A118-S132)) in the WT and mutant PrP_107-143_ amyloids. The dashed line is merely a guide for the eye. (B) Bar plot of the difference between the final C*_α_*-C*_α_* distances, *δ*C*_α_*, for *d*C*_α_*(A118-S132) and *d*C*_α_*(A118-A/V133) in the WT and mutant PrP_107-143_ amyloids. The bars and error bars represent the mean standard error (SE) of values obtained from five independent MD runs. Bars sharing the same letter are not significantly different according to Tukey’s test with *α* = 0.05. (C) Identical to (A), except for *d*C*_α_*(A120-M/V129) and *d*C*_α_*(A118-S132). Note that the *d*C*_α_*(A120-M/V129) values of PrP_107-143_ with M129V mutations (green and blue squares) are shorter than 9 Å in most of the MD runs; and their *d*C*_α_*(A120-M/V129) and *d*C*_α_*(A118-S132) values also tend to be shorter than those of the other mutants. (D) Identical to (B), except for the difference between *d*C*_α_*(A120-M/V129) and *d*C*_α_*(V122-M/V129). (E) Final snapshots of the chains D–G of PrP_107-143_(WT) and chains E–G of PrP_107-143_(M129V&I138M) after 400-ns MD simulations. The upper table shows *d*C*_α_*between the residues averaged over chains B–I. The insets show magnified views with the side chains of the residues involved in hydrophobic interactions. (F) A schematic illustration of the positional relationships and “upward shift” of the *β*-sheet(128–133) induced by the M129V mutation. The dashed lines indicate distances between the C*_α_* atoms of A118 and S132.

### 2.5 Influence of I138M mutation on the PrP_107-143_ amyloid with or without M129V mutation

Residue 138 is one of the residues that are often varied between species. For example, this residue is isoleucine in humans, methionine in many rodents, and leucine in many ruminants. Moreover, residue 138 is the most influential on the mouse–human species barrier in the transmission of sporadic CJD^25^ and also in the cross-seeding of Y145Stop peptides of human and mouse PrP.^24^ We assessed how the I138M mutation affected the local structural model of PrP^Sc^.

In a comparison between the I138M mutants and the WT, the RMSF, the *β*-sheet propensity, and the proximity scores were similar, but interestingly, the proximity scores for A133-P137 were higher in the I138M mutants, although not significantly so (Figures 5, 6, and 7). As mentioned in Section 2.4, the score for A133-P137 is related to the bending at the loop(134–137), which was also observed in I138M mutants, particularly in PrP_107-143_(M129V&I138M) (Figure 4E,F; see also Figures S3D,E and S4D,E). The angle between the *β*-sheet(128–133) and the *β*-sheet(139–141) also seemed to be near-square to acute in the I138M and M129V mutants, whereas the WT or the other mutants tended to have obtuse angles (Figures 4, S3, and S4). Figure 9 shows the proximity scores for A133-P137 of each MD run. The scores of five independent MD runs showed dispersed values in each mutant. However, the scores for A133-P137 of five MD runs were higher than 200 in PrP_107-143_(M129V&I138M), whereas in the other mutants, the scores for A133-P137 of zero to three runs were higher than 200. In the runs with relatively high proximity scores for A133-P137, the amyloids tended to form small loops like an Ω-shape (Figure 4F). Each of these loops encompassed A133 and P137, and had near-square to acute angles between the *β*-sheet(128–133) and the *β*-sheet(139–141), irrespective of the primary structures (see the boxed snapshots in Figures S3 and S4). For example, the final snapshot of Run 1 of PrP_107-143_(M129V) (Figures S3C and S4C), which had an Ω-shaped loop(134–137), also had a high proximity score (541.2; Figure 9).

**Figure 9:**
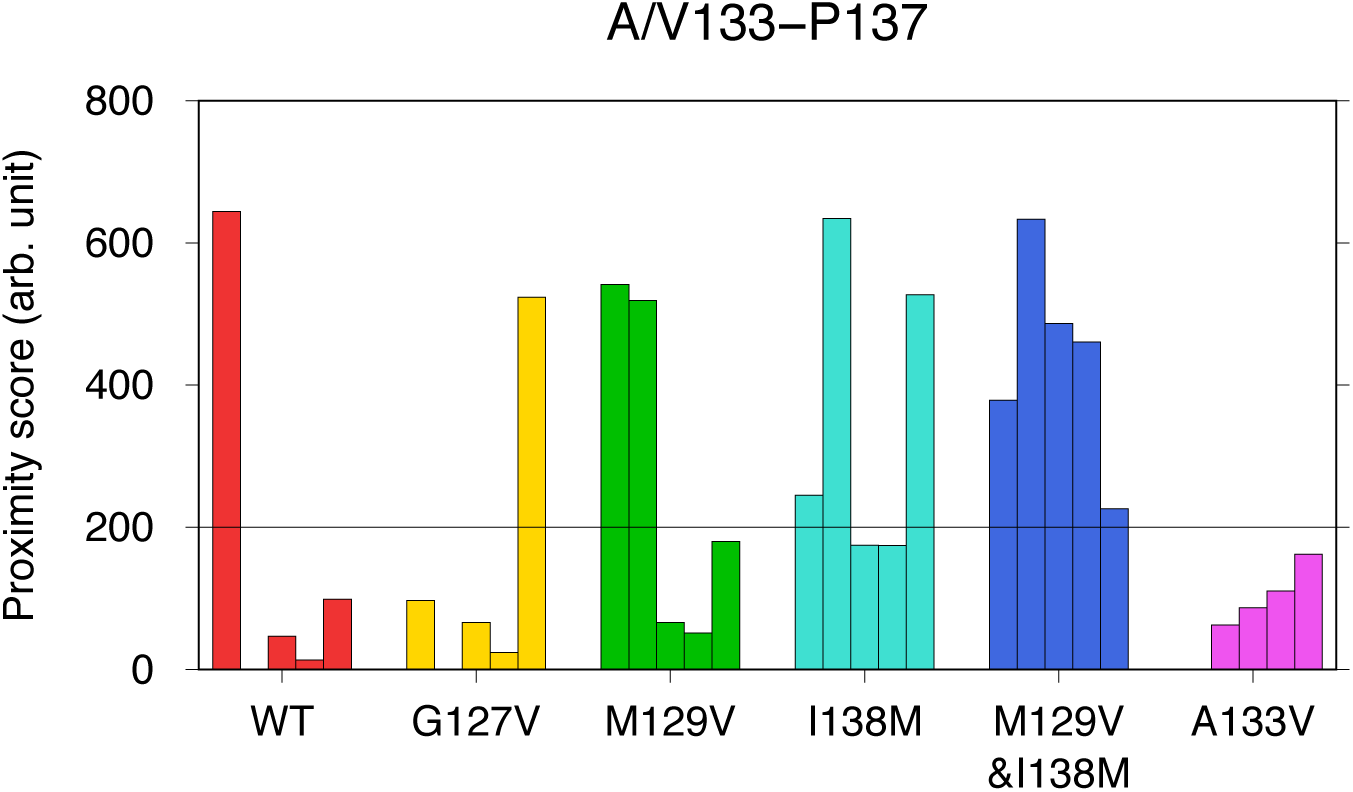
Proximity scores for A/V133-P137. Each bar represents the value obtained from one of the five runs (Runs 1 to 5, from left to right). The line at 200 is merely a guide for the eye.

The deeper bending of the loop, even to acute angles, caused by the I138M mutation was considered to be attributable to the high intrinsic *α*-helix propensity of methionine,^43, 44^ which increased the freedom of motion of the loop and the thermodynamically favorable contact between A133 and P137. In this particular conformation, V129 seemed to be compatible with M138; that is, the “upward shift” of the *β*-sheet(128–133) brought A133 into sufficient proximity with P137 for their efficient interaction, and allowed loop(134–137) to bend deeper, without the C-terminal-side region colliding against loop(116–118) (Figure 8F). As a result, the interaction of A118-P137 was considerably weakened in the M129V mutants (Figure 7). In contrast, in the mutants with M129, the full degree of deeper bending was not allowed due to steric clash. This may explain why the effects of the I138M mutation were more accentuated in PrP_107-143_(M129V&I138M) than in PrP_107-143_(I138M). For example, the proximity score of V122-L125 in PrP_107-143_(M129V&I138M) was higher than that in PrP_107-143_(M129V) by about 200, whereas the proximity scores in PrP_107-143_(WT) and PrP_107-143_(I138M) were almost equal (Figure 6). It is possible that the efficient bending of loop(134–137) in PrP_107-143_(M129V&I138M) facilitated further shifting of the *β*-sheet(128– 133) to cause the difference, whereas the bending tended to be hampered by M129 in PrP_107-143_(I138M) and the effects of M138 could not be fully manifested. The same explanation holds for the different proximity scores for A116-I139, A117-A133, and A118-A133 (Figure 7).

### 2.6 Influence of A133V mutation on the PrP_107-143_ amyloid

V136 (in ovine numbering) is one of the scrapie-prone polymorphisms of ovine PrP. ^45^ In the present model shown in Figure 4A, the corresponding residue of human PrP, A133, faced the same direction as M129; we were therefore interested in whether an A133V mutation would somehow affect the PrP_107-143_ amyloid structures. The mutant PrP_107-143_(A133V) appeared to have a *β*-sheet(120–122) and *β*-sheet(128–133) that were slightly more stable than those of the WT overall (Figures 4G, S3F, and S4F). In the heatmap patterns, these regions were characterized by more demarcated *β*-sheet propensities and lower RMSF values (Figure 5). Consistent with this appearance, the distances between the C*_α_* atoms of residues 118 and 133 were very stably maintained at *∼* 7 Å (Figure 10), with the proximity score for A118-V133 being significantly higher than that for the WT (Figure 7). The interaction of A118-V133 may cause a significant increase in the proximity scores for L130-L130 and M134-M134. In the U-shaped loop, the score for V122-M129 was significantly higher and, reciprocally, those for A120-M129 and V122-L125 were slightly lower in the A133V mutant than in the WT, although these decreases were not statistically significant (Figure 6). The hydrophobic interaction in the N-/C-terminal-side regions was also affected: the score for A117-I139 was increased, although not significantly so. This increase could be explained from the viewpoint of the positional relationship between *β*-sheet(120–122) and *β*-sheet(128–133), as discussed above (Figure 8E,F). In the A133V mutant, the stable A118-V133 relationship and the stacking interactions of L130 and M134 would fix the positional relationship of the two *β*-sheets as in the left panel of Figure 8F. This positional relationship would facilitate interactions of V122-M129 and A117-P139, because of the restricted motion range of the C-terminal-side region.

**Figure 10:**
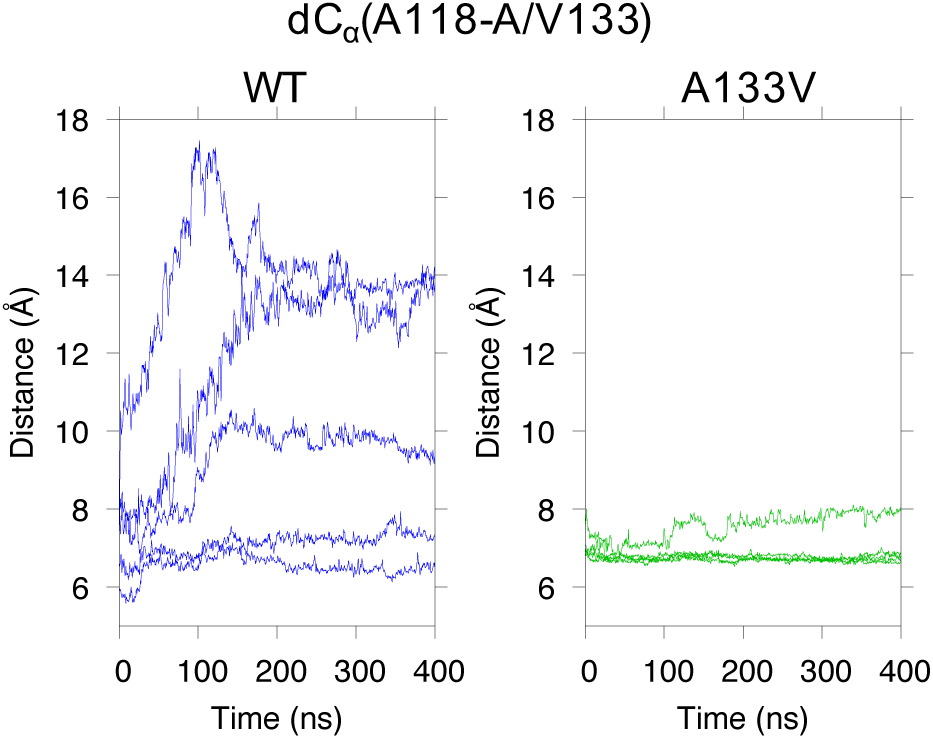
Fluctuations in the average distance between C*_α_* atoms of A118 and A/V133, *d*C*_α_*(A118-A/V133). Each line represents the average of the distances over chains B–I in each of the five MD runs.

The altered balance of the hydrophobic interactions induced by the A133V mutation did not improve the stability of the amyloid as much as the M129V mutation, despite the stabilized relationships of the two *β*-sheets; thus, the simulations of the present model did not provide any clues to the mechanism by which V136 of ovine PrP facilitates conversion to disease.

### 2.7 Conclusions about the PrP_107-143_ amyloid model

Collectively, the above results suggested that the present local structural model of PrP^Sc^ was more compatible with V129 than with M129. This is an example of an amyloid conformation in which differences between methionine and valine affect the structural stability. Another model that is more compatible with M129 than with V129 is underway.

Reinforcement of a single hydrophobic interaction between the *β*-sheets induced by mutations, for example, the V122-M129 interaction in PrP_107-143_(A133V) or A120-V129 interaction in PrP_107-143_(M129V&I138M) amyloids, did not facilitate stabilization of the local structures. To the contrary, the significantly lowered L125-L125 proximity score of PrP_107-143_(M129V&I138M) suggested that this alteration caused a certain degree of destabilization of the *β*-arch (Figures S3E and S4E). The results of *α*Syn(G84M) and PrP_107-143_(M129V) implied that well-balanced interactions of constituent residues in the hydrophobic core might be essential for the maximal stability of a *β*-arch.

Incidentally, the spontaneous approach of the N- and C-termini of the PrP_107-143_ amyloid, which occurred with high A/V133-P137 proximity scores, was notable because the N- and C-termini led to positive- and negative-charge clusters, respectively, in a full-length human PrP. Those charged clusters were able to electrostatically interact and contribute to the stabilization of the amyloid structures. Interestingly, the positive-charge cluster was previously suggested to be essential for the structural conversion of the N-terminal region. ^46^

Knowledge from the present study may help in the design of useful amyloids or prediction of potentially-amyloidogenic proteins.

### 2.8 Generally applicable findings of the present study

The unique effects of each group of hydrophobic amino acids in the in-register parallel *β*-sheet amyloids were conformation-dependent and attributable to C*_β_*-branching and/or the length of the side chains. Methionine and isoleucine showed similar influences in the flat-loop of the *α*Syn amyloid (E61M and E61I), whereas isoleucine and valine in the U-shaped loops destabilized the local structures when they were not stably incorporated in hydrophobic cores (G84I and G84V). As the latter effects were observed in two different protein models, *α*Syn(G84I) and PrP_107-143_(G127V), this notion may be generally applicable to other in-register parallel amyloids. In certain situations, methionine can behave distinctly from other hydrophobic amino acids, as in *α*Syn(G84M) amyloid, where its long side chain functions as a crossbeam that runs across the *β*-arch and stabilizes it, forming a well-balanced interaction network. The long hydrophobic side chain of methionine would also be advantageous for efficient inter-domain and intermolecular interactions, as seen in an amyloid *β* (A*β*) fibril (PDB ID: 2MPZ).^47^ However, methionine is not always advantageous for stable hydrophobic core formation, as demonstrated in the M129V mutants of the PrP_107-143_ model amyloid. In terms of C*_β_*-branching and the length of side chains, valine and methionine can have mutually different influences on *β*-arches, and, if the *β*-arch constitutes an interface between the substrate monomer and template amyloid, such differences would cause species/strain barriers. The influences of an M/V129 polymorphism of human PrP on the phenotypes of prion diseases and susceptibilities to certain prion strains, for example, new-variant CJD, ^7, 8^ might be explainable from this view point.

As mentioned in Section 1, while our study was under revision, the cryo-EM structures of brain-derived prion proteins (263K and GPI-anchored/anchorless RML strains) were reported.^20–22^ Figure 11 shows the cryo-EM structure of the 263K prion protein (PDB: 7LNA^20^), which has an in-register parallel *β*-sheet structure with a Greek-key motif at its hydrophobic core, residues 112–134. This feature fits nicely with our model, but some differences are also apparent between the two models. In our model, for example, the side chain of M129 points inward of the U-shaped loop (Figure 4A), whereas it points outward in the cryo-EM structure. Moreover, these structures have different hydrophobic interaction networks in the Greek-key domain. Further investigation is needed to elucidate the effect of mutation (including M129V) on the 263K strain.

**Figure 11:**
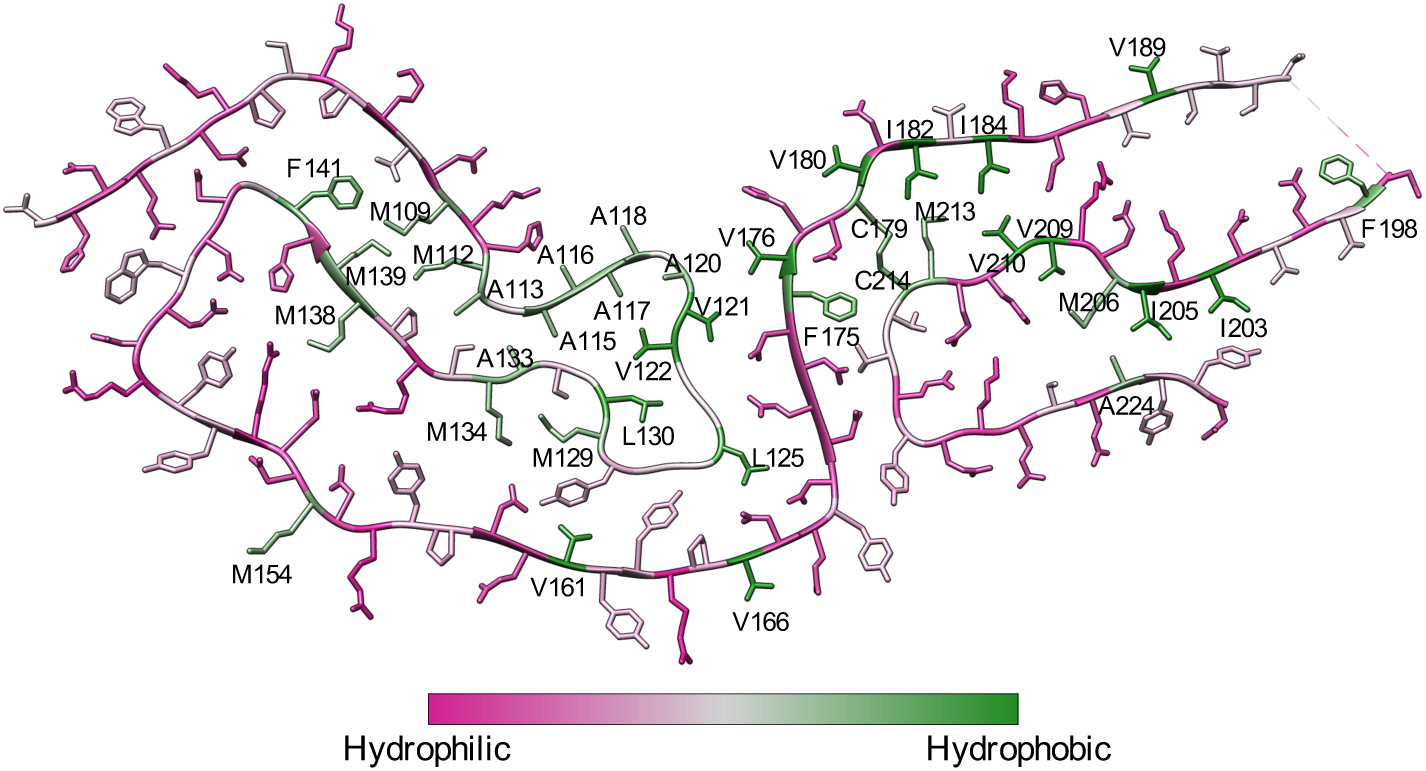
Cryo-EM structure of a brain-derived 263K prion protein (PDB: 7LNA^20^). Residues are colored using a Kyte-Doolittle hydrophobicity scale, ^87^ and hydrophobic residues are labeled.

Additionally, the cryo-EM structure has other hydrophobic cores around M109, M206, and M213. Different hydrophobic amino acids at codons 109 and 112 influence the transmission efficiencies of prions among different hamster species.^26, 27^ In an experiment observing the influences of systematically introduced mutations of mouse PrP on the conversion efficiencies in scrapie-infected Neuro2a cells, methionine at some positions (e.g., M213) was shown to be replaceable with other hydrophobic residues such as leucine without affecting the conversion efficiency, whereas it was irreplaceable at other residues (e.g., M206). ^48^ The results of the present study might assist in the analysis of these hydrophobic interactions to find clues explaining the transmission efficiencies, and might be applicable to other amyloids.

In hamster 263K PrP^Sc^, the C-terminus and a portion of the third *α*-helix (residues 199– 227) form a pair of *β*-sheets with a 180-degree turn, that is, *β*-arch, in between. Interestingly, the *β*-arch contains residue 219 whose E/K polymorphism of human PrP reportedly greatly affect susceptibility to sporadic CJD, ^49–51^ while not to variant CJD (vCJD).^52, 53^ Our previous experiments with cultured cells exhibited that a mutant PrP with a disulfide crosslink between residues 168 and 225 (in mouse numbering) was converted to protease-resistant form by RML or 22L mouse prions but not by Fukuoka-1 mouse-adapted Gerstmann-Sträussler-Sheinker syndrome (GSS) prion. ^54^ This suggested that the position of the C-terminus of GSS PrP^Sc^ was different from that of RML PrP^Sc^. If residue 219 of human PrP is influential on the stability of the *β*-arch comprising it, it may explain why CJD and GSS (or vCJD) are differently affected by E/K219 polymorphism. In the case, stability of *β*-arches is thus important for strain diversity of prions.

### 2.9 Limitations to the present method

A caveat to our hypothesis is that the MD simulation is an artificial setting, which starts with the same initial conformations irrespective of mutant types. In reality, distinct conformations are implied for some mutants. For example, the spontaneous amyloid formation caused by the Y145Stop mutant of Syrian-hamster PrP, which has more methionine than the mouse or human PrP counterparts, is inefficient.^27^ We consider that the higher freedom of motion of the backbone and potential steric effects of the long side chain of methionine hamper its settling into amyloid conformations. Such inefficiency in folding into a conformation may divert a certain fraction of molecules to aberrant refolding pathways in in vitro or in vivo experiments. Regardless of these limitations, MD simulation provides direct insight into the structures and dynamics of in-register parallel *β*-sheet amyloids, and this approach would shed light on the mechanisms of the strain diversity of amyloids, including PrP^Sc^. One approach to overcome this limitation is generating initial structures for all-atom MD simulations with coarse-grained (CG) MD simulations and backmapping. ^55^ The CG models allow us to simulate the dynamics of biomolecular systems on a longer timescale; thus, the effect of a mutation might be reflected after a longer-term CG MD simulation.

Another caveat concerns the force field for the MD simulations. Many studies have assessed the quality of the force fields for amyloidogenic proteins using a segment or the full length of A*β* peptides. ^56–63^ In this study, we used the AMBER ff99SB-ILDN force field, ^64^ which provides fairly good performance and is one of the safest choices for this purpose. ^57, 58, 60^

However, Watts et al. reported that several force fields including AMBER ff99SB-ILDN yield a positive value of binding energy in the A*β*(1–40) dimer, resulting in instability of the quaternary structures. ^59^ In contrast, CHARMM36 shows favorable protein–protein interactions.^59^ The CHARMM36 force field was refined for the simulation of intrinsically disordered proteins (CHARMM36m).^65^ Samantray et al. demonstrated that CHARMM36m is suitable for simulating peptide aggregation. ^56^ From these points, we expect that the MD simulation with CHARMM36(m) can stabilize our systems (i.e., *α*Syn and PrP_107-143_) and highlight the instability caused by the mutation.

## 3 Conclusion

We have demonstrated how different hydrophobic amino acids uniquely affect the stability of U-shaped *β*-arches and consequently the entire amyloid in two different in-register parallel *β*-sheet amyloids. Our studies have also revealed how the manifestations of one mutation are affected by another mutation. This concept would be applicable to various in-register parallel *β*-sheet amyloids and other *β*-arches of PrP^Sc^. We expect that the knowledge from the present study will contribute to prediction of potentially-amyloidogenic proteins or proteins which might interact with a given pathogenic amyloid in the future, in addition to advancing our understanding of the strain diversity of amyloids.

## 4 Methods

### 4.1 Modeling structures of ***α***Syn amyloids

We used a Greek-key *α*Syn amyloid (PDB ID: 2N0A^34^) as a starting structure, after truncating the disordered N- and C-terminal-side regions (residues 1–35 and 100–140, respectively) (see Figure 1).^30^ The N- and C-termini were acetylated and N-methylated using Amber-Tools16, respectively. ^66^ For modeling the *α*Syn mutants, we used SCWRL4.^67^ The modeled amyloids were solvated with a rhombic dodecahedron water box with a minimum wall distance of 12 Å using GROMACS (version 5.0.4). Na^+^ and Cl*^−^* ions were randomly placed to neutralize the system and yield a net concentration of 150 mM NaCl. The protonation state of the histidine at residue 50 was fixed as the N*_δ_*-H tautomer (HID form in the AMBER naming convention) in all simulations.

### 4.2 MD simulation of the ***α***Syn amyloids

GROMACS (versions 5.0.4 and 5.1.2) ^68^ with the AMBER ff99SB-ILDN force field ^64^ was used for MD simulations with the TIP3P water model.^69^ The system was initially minimized for 5,000 steps with the steepest descent method, followed by 2,000 steps with the conjugate gradient method. During minimization, heavy atoms of the amyloids were restrained with a harmonic potential with a force constant of 10.0 kcal*/*mol *·* Å^2^. After minimization, the system temperature was increased from 0 to 310 K during a 1 ns simulation with the restraints. Next, a 1 ns equilibration run was performed while gradually removing the restraints from 10.0 kcal*/*mol *·* Å^2^ to zero, and subsequent equilibration was performed in the NPT ensemble for 2 ns at 310 K and 1 bar. The production runs were carried out for 400 ns in the NPT ensemble at 310 K and 1 bar (Figure S7A). We used the velocity-rescaling scheme^70^ and the Berendsen scheme^71^ for the thermostat and barostat, respectively. The LINCS algorithm^72^ was used to constrain all bonds with hydrogen atoms, allowing the use of a 2 fs time-step. Electrostatic interactions were calculated with the Particle-mesh Ewald method. ^73^ The cut-off length was set to 12 Å for the Coulomb and van der Waals interactions. The Verlet cutoff scheme^74^ was used for neighbor-searching. Trajectory snapshots were saved every 10 ps. We conducted five (for WT) and three (for the mutants) independent 400-ns MD simulations.

### 4.3 Analyses

Backbone root-mean-square deviation (RMSD), RMSF, potential energy, and distance between atoms were calculated using GROMACS. ^68^ Convergence of the MD simulations was assessed with the root-mean-square inner product (RMSIP)^75^ between two halves of the last 300 ns of the trajectories. The essential subspace was extracted by using principal component analysis (PCA) for C*_α_* atoms. Table S1 shows that the first 20 PCs explain 69% or more of the variance for all the simulations of *α*Syn. From this result, the first 20 eigenvectors of C*_α_* atoms were used to calculate RMSIP. The RMSIP values are also summarized in Table S1. In all the simulations of *α*Syn, the RMSIP values are larger than 0.6, which suggests good convergence of the trajectories. ^75^ Backbone RMSD (Figure S8) and potential energy (Figure S9) support the convergence of this region.

The secondary structure content during the simulations was calculated with DSSP ^76, 77^ using *gmx do dssp* in GROMACS. Hydrophobic contacts were analyzed using PyInteraph. ^78^ A hydrophobic contact was assigned if the distance between the centers of mass of the side chains was less than 5 Å (Figure S6A).^42, 78^ The results of the hydrophobic contact analyses were visualized using Cytoscape (version 3.5.1). ^79^ All molecular structures for the amyloids were drawn with UCSF Chimera (version 1.12) ^80^ or VMD.^81^ Movies of the MD simulation trajectories were prepared with UCSF Chimera. ^80^

### 4.4 Modeling PrP_107-143_ amyloids and MD simulations

In designing the local structural models of human PrP^Sc^, we adopted the region comprising *−*GGL_125_GG*−*, as denoted above (see Section 2.2), and avoided charged residues in the N- and C-termini of the peptide because they can excessively destabilize in-register parallel *β*-sheet structures, particularly when they are at the free ends. There were positive- and negative-charge clusters in regions 101–106 and 144–147, respectively; thus, the region between them was used for the modeling. First, we roughly designed conformations of a single layer of amyloid with Swiss PDB viewer^82^ based on the structural model proposed by Theint et al., ^39^ then piled it up at intervals of about 5 Å to make an in-register parallel *β*-sheet amyloid with UCSF Chimera. ^80^ The model amyloids with 10 layers were refined using Modeller (version 9.15)^83^ with *β*-sheet restraints for the residues A120–V122, Y128–L130, and I138–F141. Mutants were generated by using SCWRL4,^67^ and subsequently the N- and C-termini of the refined models were acetylated and N-methylated using PyMOL, respectively. ^84^ We performed five independent 400-ns MD simulations for each model using nearly the same procedure as for the MD simulations of *α*Syn (Figure S7B). We checked the convergence of the simulations with RMSIP (Table S2), backbone RMSD (Figure S10), and potential energy (Figure S11). Table S2 summarizes the RMSIP between two halves of the last 100 ns of the trajectories and cumulative proportion variance evaluated with the first 20 PCs. The RMSIP values are larger than 0.6 in all the simulations, and the first 20 PCs explain over 65% of variance in most cases. We thus used the last 100 ns for the analyses described in Section 4.3.

### 4.5 Statistical analyses

Tukey’s multiple comparison test (*α* = 0.05) was applied for the statistical analysis with the aid of the *multcomp* package in R.^85^ The graphs of the statistical analysis were drawn with the *ggplot2* R package.^86^

## Associated Content

### Supporting Information

The Supporting Information is available free of charge at ***.

• SI.pdf: Radius of gyration as a function of time for the *α*Syn mutants; C*_α_*-C*_α_* distance between residues 84 and 89 in the *α*Syn mutants; final snapshots of MD simulations of PrP_107-143_; radius of gyration as a function of time for the PrP_107-143_; definition of hydrophobic contact and proximity score; correlations between proximity score and C*_α_*-C*_α_*distance; procedure of MD simulation; RMSIP and cumulative proportion of variance of *α*Syn; time series of backbone RMSD of the *α*Syn amyloids; time series of potential energy of *α*Syn amyloids; RMSIP and cumulative proportion of variance of PrP_107-143_; time series of backbone RMSD of PrP_107-143_; time series of potential energy of PrP_107-143_.

• Movie S1.mpg: A trajectory of MD simulation of *α*Syn(G84M). Only residues 82 to 91 are shown. The side chains of M84 and A89 are shown with sticks.

• Movie S2.mpg: A trajectory of MD simulation of PrP_107-143_(WT).

• Movie S3.mpg: A trajectory of MD simulation of PrP_107-143_(G127V).

## Acknowledgments

The numerical calculations were performed on the TSUBAME2.5/3.0 supercomputer at the Tokyo Institute of Technology and the Reedbush supercomputers at the Information Technology Center, the University of Tokyo. HO acknowledges the support of the “TSUBAME Encouragement Program for Young/Female Users” of the Global Scientific Information and Computing Center at the Tokyo Institute of Technology; and the “Initiative on Promotion of Supercomputing for Young or Women Researchers” of the Information Technology Center, the University of Tokyo. This work was supported by a grant from the Takeda Science Foundation (www.takeda-sci.or.jp/) (to YT), and JSPS KAKENHI Grant Numbers 19K16058 (to HO) and 19K22539 (to YT).

## Author Contributions

Conceptualization, Y.T.; Methodology, H.O.; Validation, Y.T. and N.N.; Formal Analysis, Y.T. and H.O.; Investigation, H.O. and Y.T.; Resources, H.O. and N.N.; Data Curation, H.O. and Y.T.; Writing (Original Draft Preparation), Y.T.; Writing (Review & Editing), Y.T., H.O., and N.N.; Visualization, H.O. and Y.T.; Supervision, N.N.; Project Administration, Y.T.; Funding Acquisition, H.O., Y.T., and N.N.

## Notes

The authors declare no competing financial interests.

