## Supporting Information for "Conformation-dependent influences of hydrophobic amino acids in two in-register parallel *β*-sheet amyloids, an *α*-synuclein amyloid and a local structural model of PrP^Sc^"

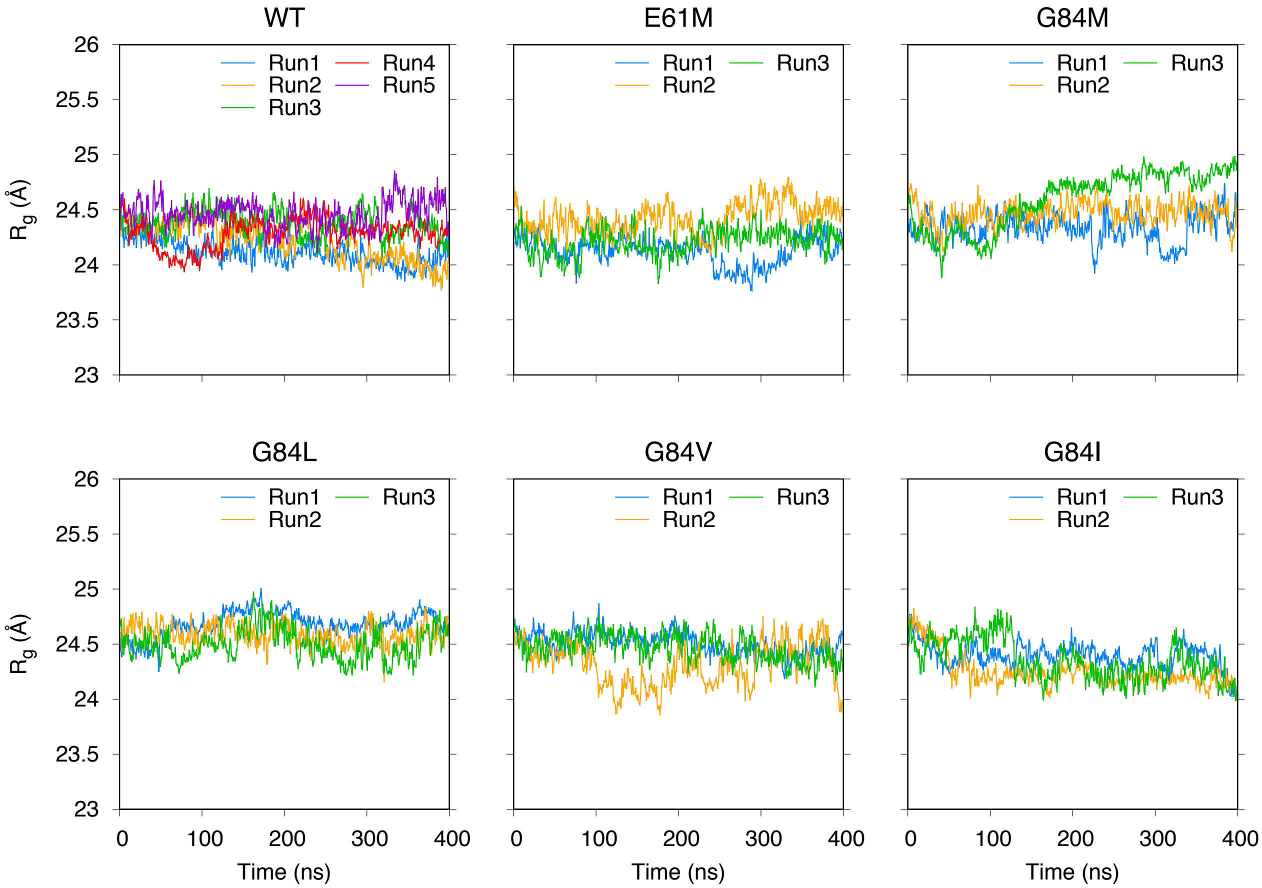


Figure S1: Radius of gyration ($R_g$) as a function of time for the $\alpha$Syn mutants.

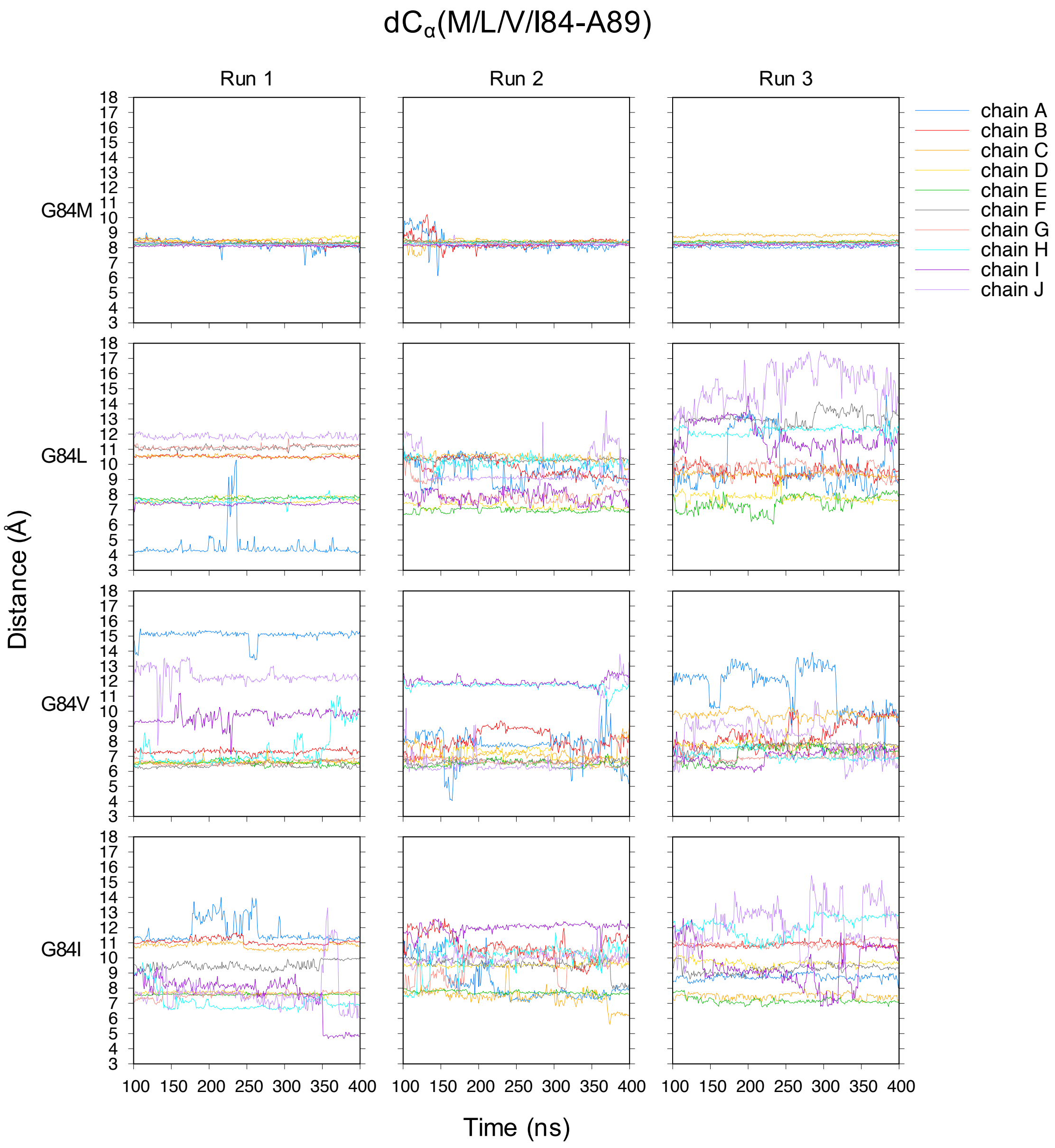


Figure S2: Time series of distance between $C_\alpha$ atoms ($dC_\alpha$) of residues M/L/V/I84 and A89 in the $\alpha$Syn mutants.

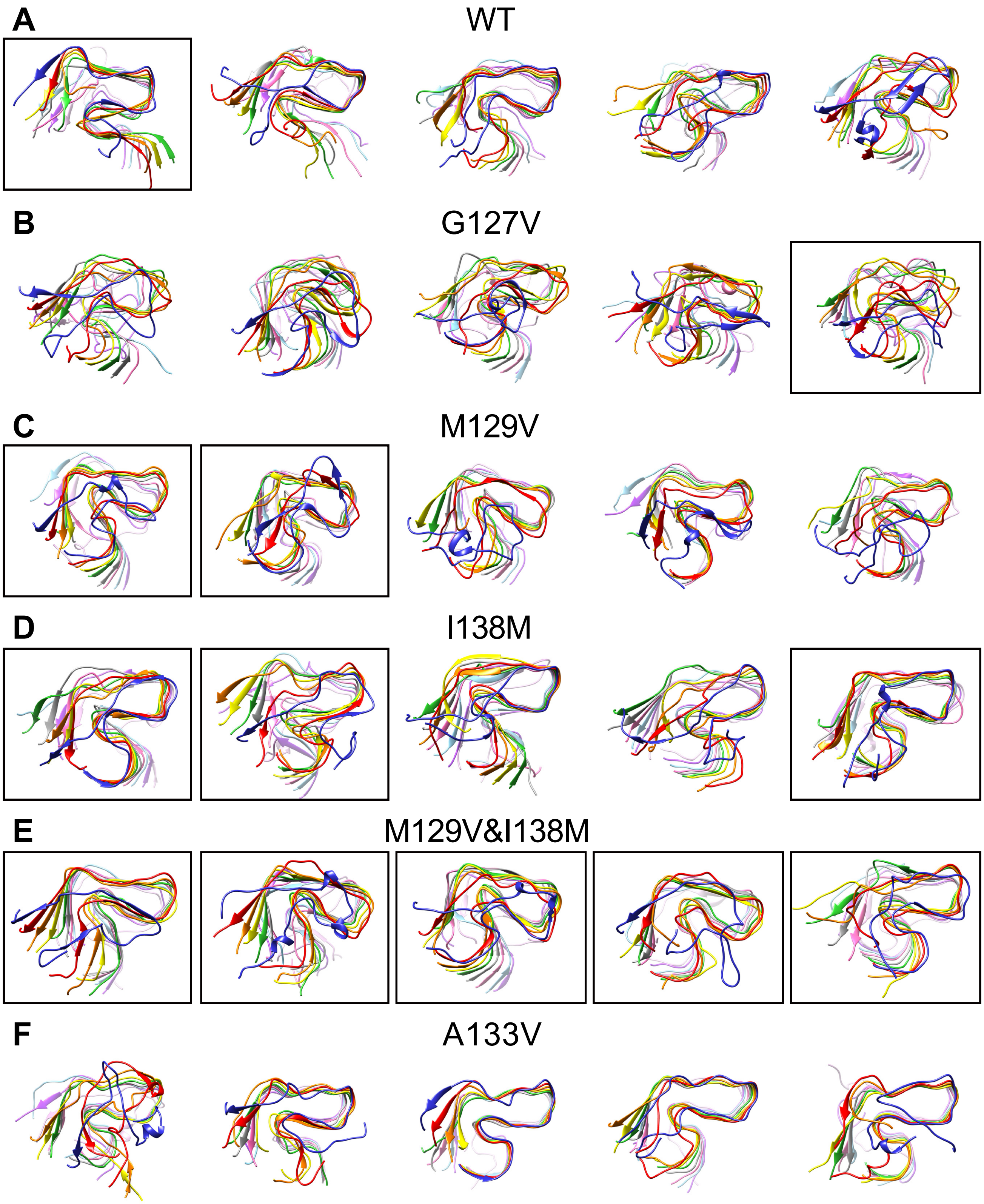


Figure S3: Final snapshots of the five independent runs of MD simulation of $PrP_{107\text{-}143}$: (A) WT, (B) G127V, (C) M129V, (D) I138M, (E) M129V&I138M, and (F) A133V. From left to right, top to bottom: Runs 1–5. Those with proximity scores for A133-P137 higher than 200 are in black boxes. See also Figure 9 in the main text.

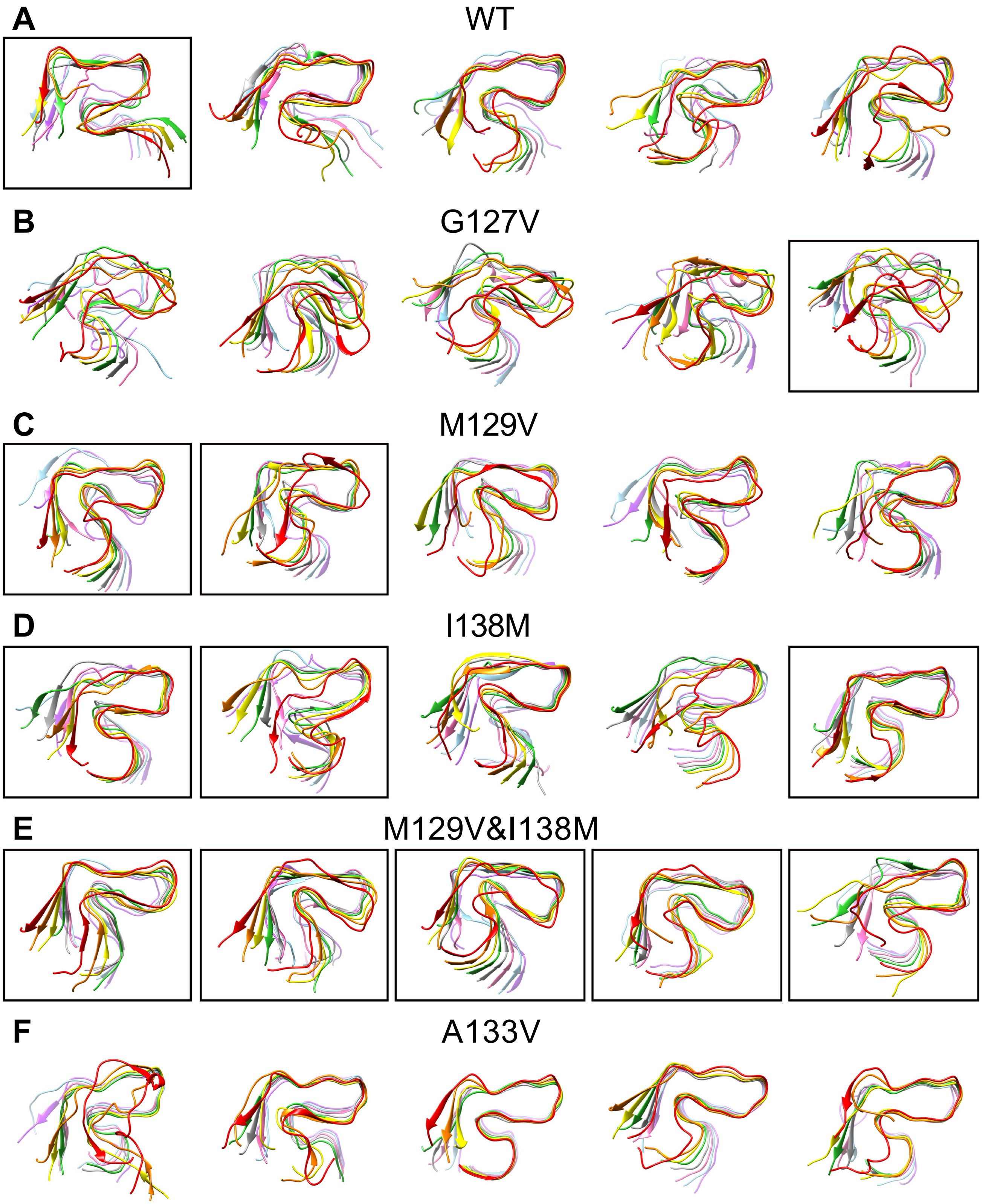


Figure S4: Identical to Figure S3, except that chains A and J are removed for clarity.

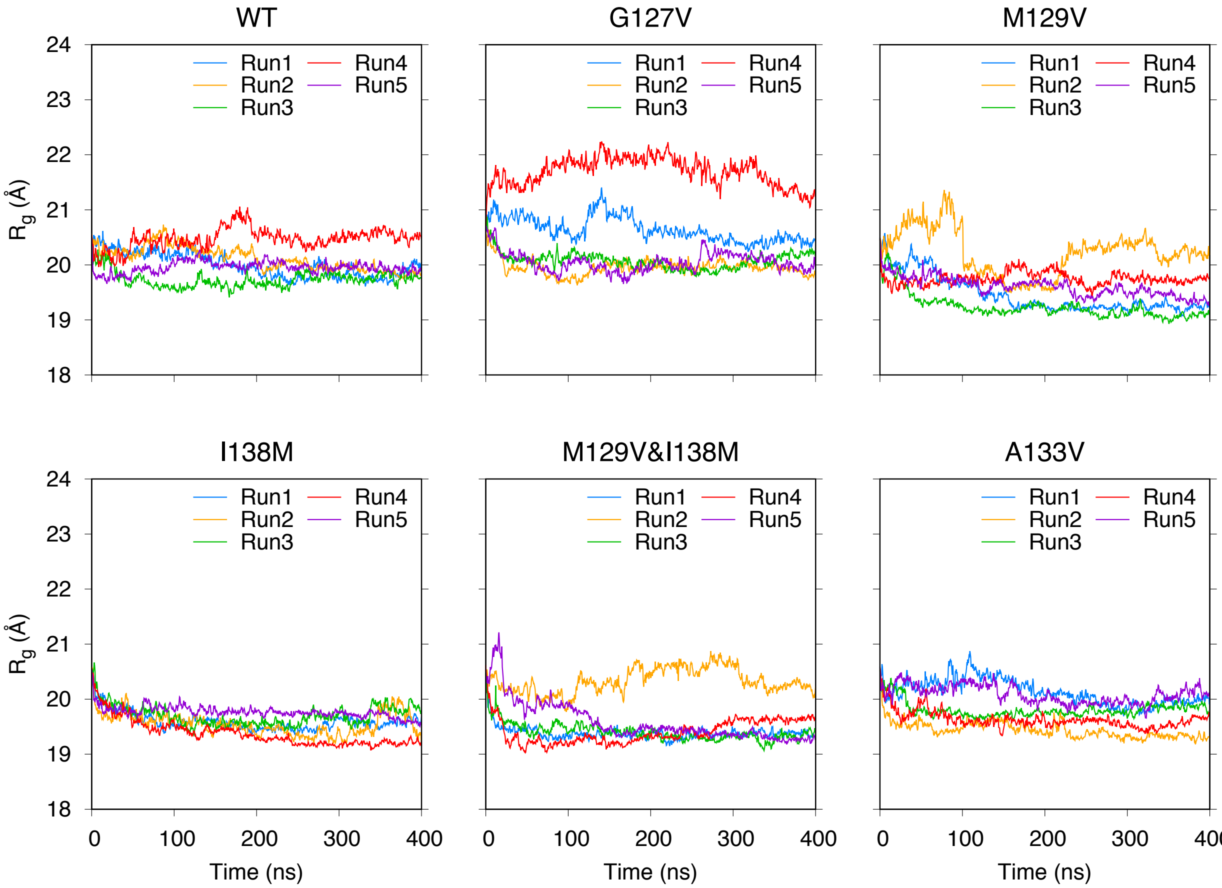


Figure S5: Identical to Figure S1, except for the $PrP_{107\text{-}143}$.

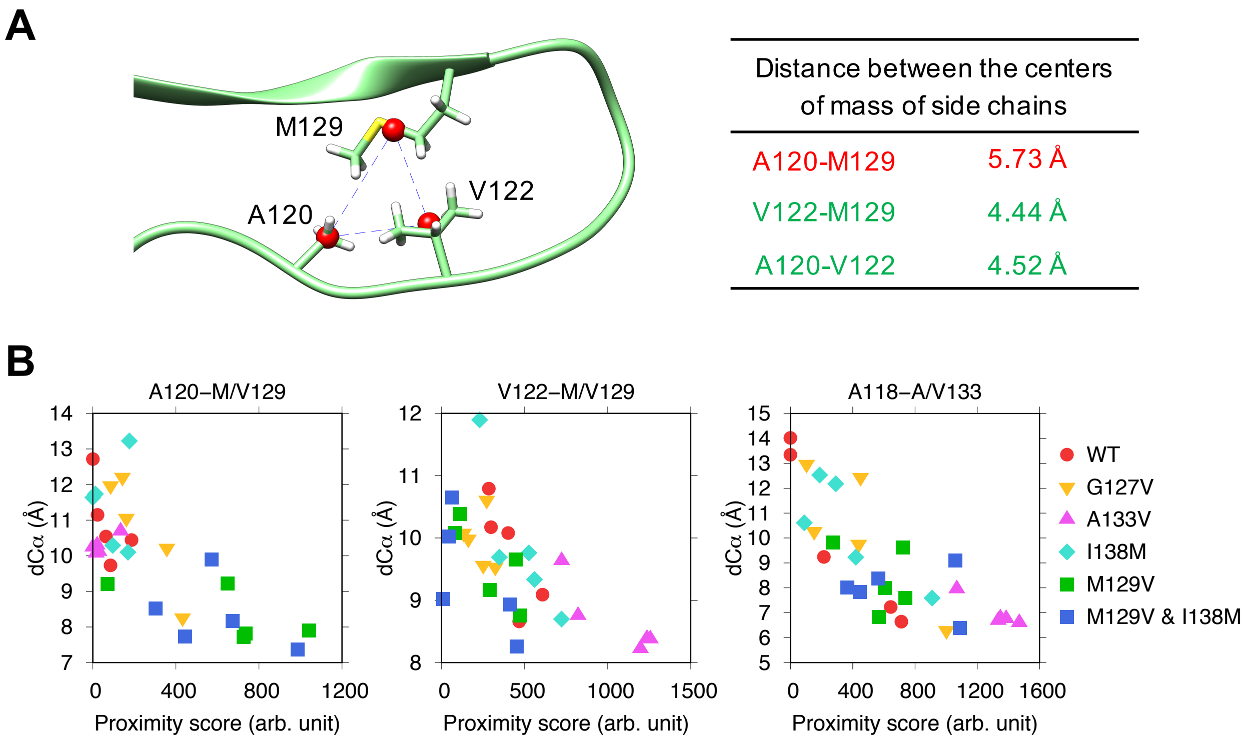


Figure S6: (A) Definition of hydrophobic contact and proximity score. A "hydrophobic contact" is counted when the distance between the centers of mass of side chains (red spheres in this figure) of two hydrophobic residues are within 5 Å.[1] For example, in this figure, V122-M129 and A120-V122 are counted as contacts but A120-M129 is not. A "proximity score" basically sums up the proportions of time/snapshots in a trajectory when hydrophobic contact is present between the two residues of interest over chains B–I. When a single residue has hydrophobic contact simultaneously with a residue on the same chain and another residue of the same residue number on the next chain, the score for the interaction in the snapshot is two. Therefore, the total proximity score can be larger than 800 if there are many inter-chain interactions. (B) Correlations between proximity score and $dC_\alpha$ between the corresponding residues averaged over eight chains in the last snapshot of MD simulation.

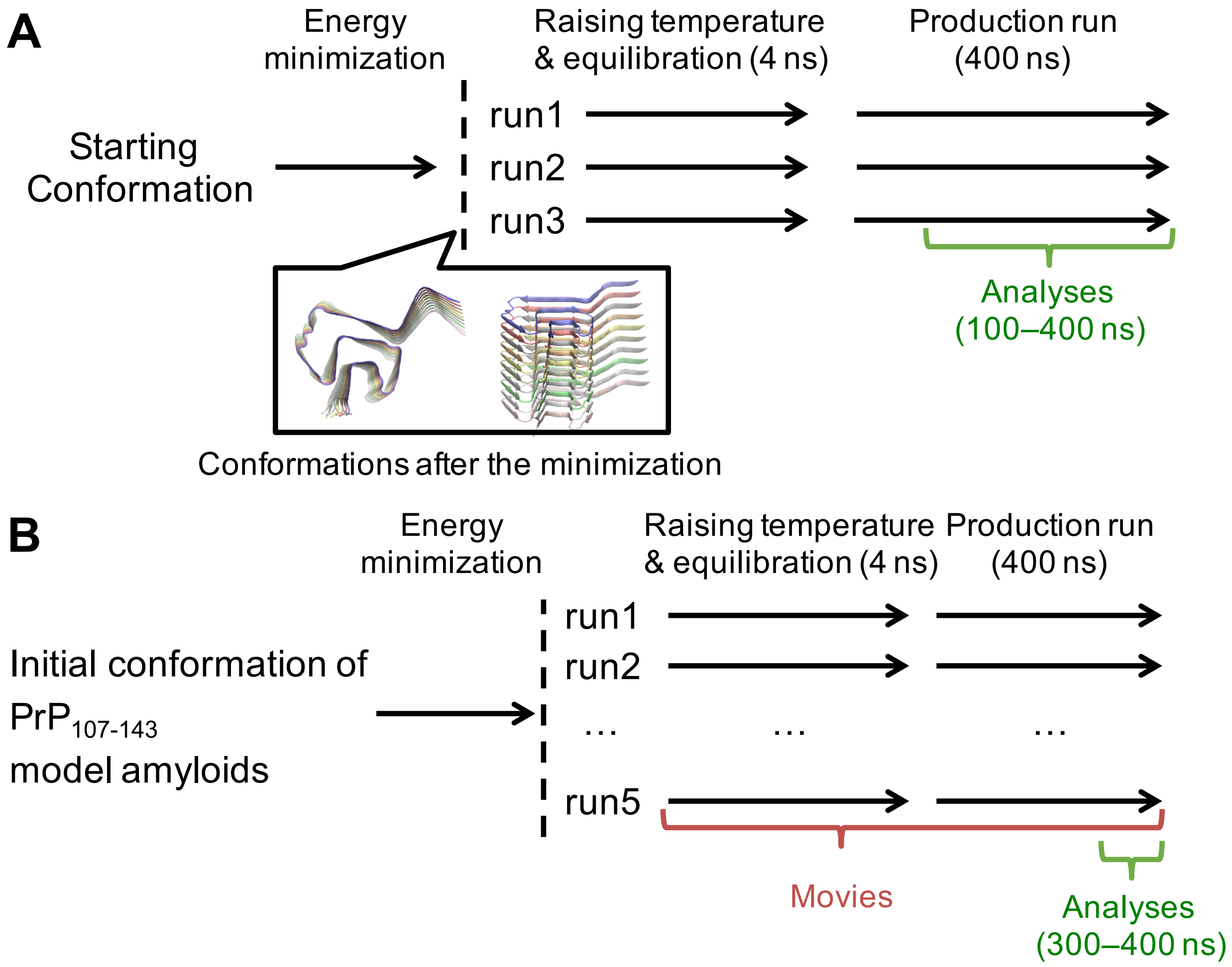


Figure S7: Procedure of MD simulation: (A) MD simulation of $\alpha$Syn. (B) MD simulation of the tentative local structural model of $PrP^{Sc}$, $PrP_{107\text{-}143}$ amyloids.

Table S1: Root-mean-square inner product (RMSIP) and cumulative proportion of variance of $\alpha$Syn calculated with the first 20 principal components

| Run | WT | | E61M | | G84M | |
|---|---|---|---|---|---|---|
| | RMSIP | Cumulative % | RMSIP | Cumulative % | RMSIP | Cumulative % |
| 1 | 0.74 | 69.4 | 0.72 | 85.4 | 0.74 | 83.3 |
| 2 | 0.70 | 74.9 | 0.69 | 75.8 | 0.75 | 77.7 |
| 3 | 0.73 | 74.8 | 0.79 | 74.4 | 0.76 | 80.8 |
| 4 | 0.72 | 74.0 | | | | |
| 5 | 0.77 | 81.7 | | | | |

| Run | G84L | | G84V | | G84I | |
|---|---|---|---|---|---|---|
| | RMSIP | Cumulative % | RMSIP | Cumulative % | RMSIP | Cumulative % |
| 1 | 0.75 | 69.7 | 0.73 | 74.0 | 0.69 | 81.7 |
| 2 | 0.68 | 81.1 | 0.69 | 81.1 | 0.65 | 75.6 |
| 3 | 0.64 | 74.7 | 0.73 | 72.0 | 0.71 | 81.0 |

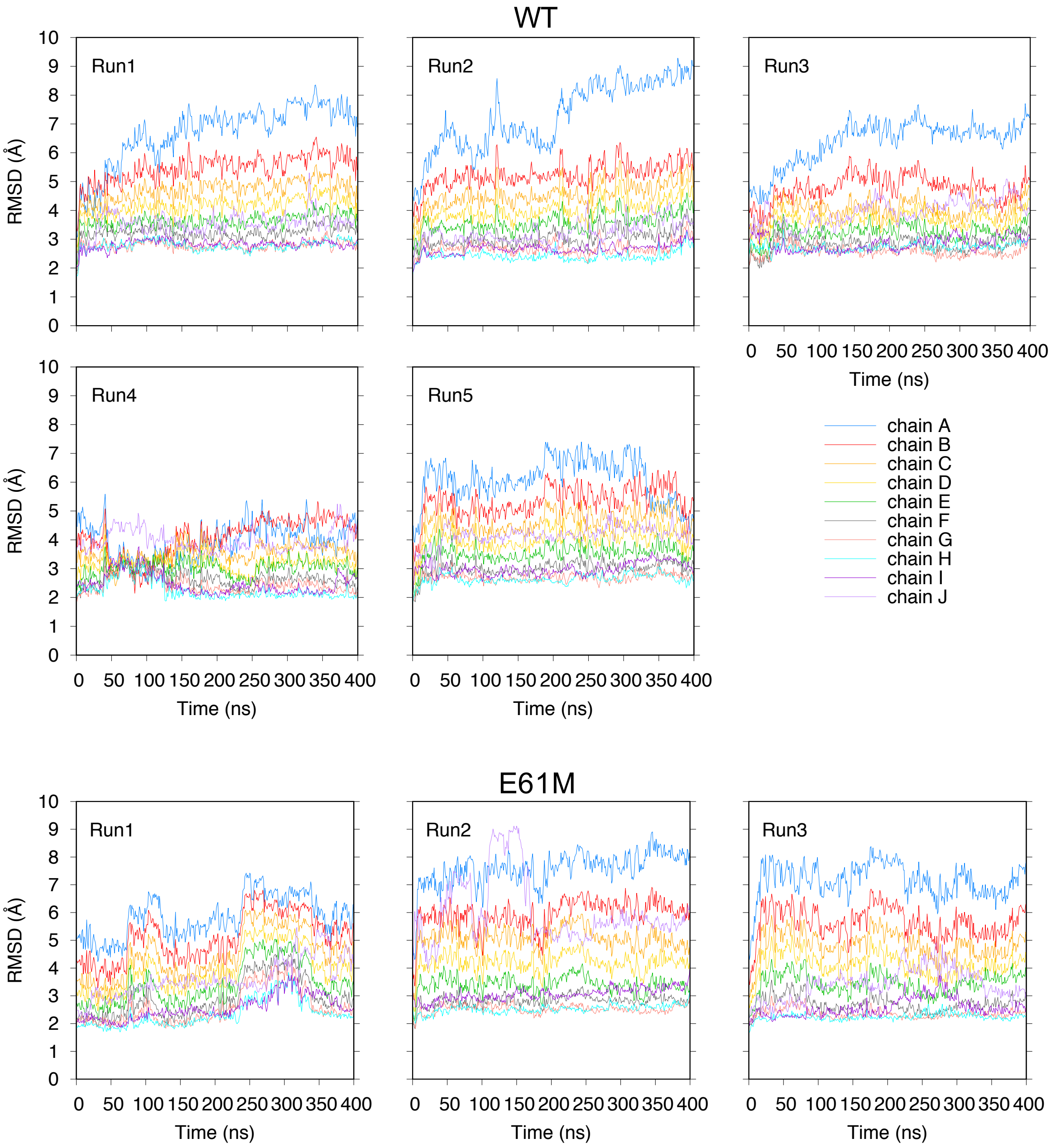


Figure S8: Backbone root-mean-square deviation (RMSD) of the $\alpha$Syn amyloids with respect to the solid-state NMR structure (PDB ID: 2N0A[2]).

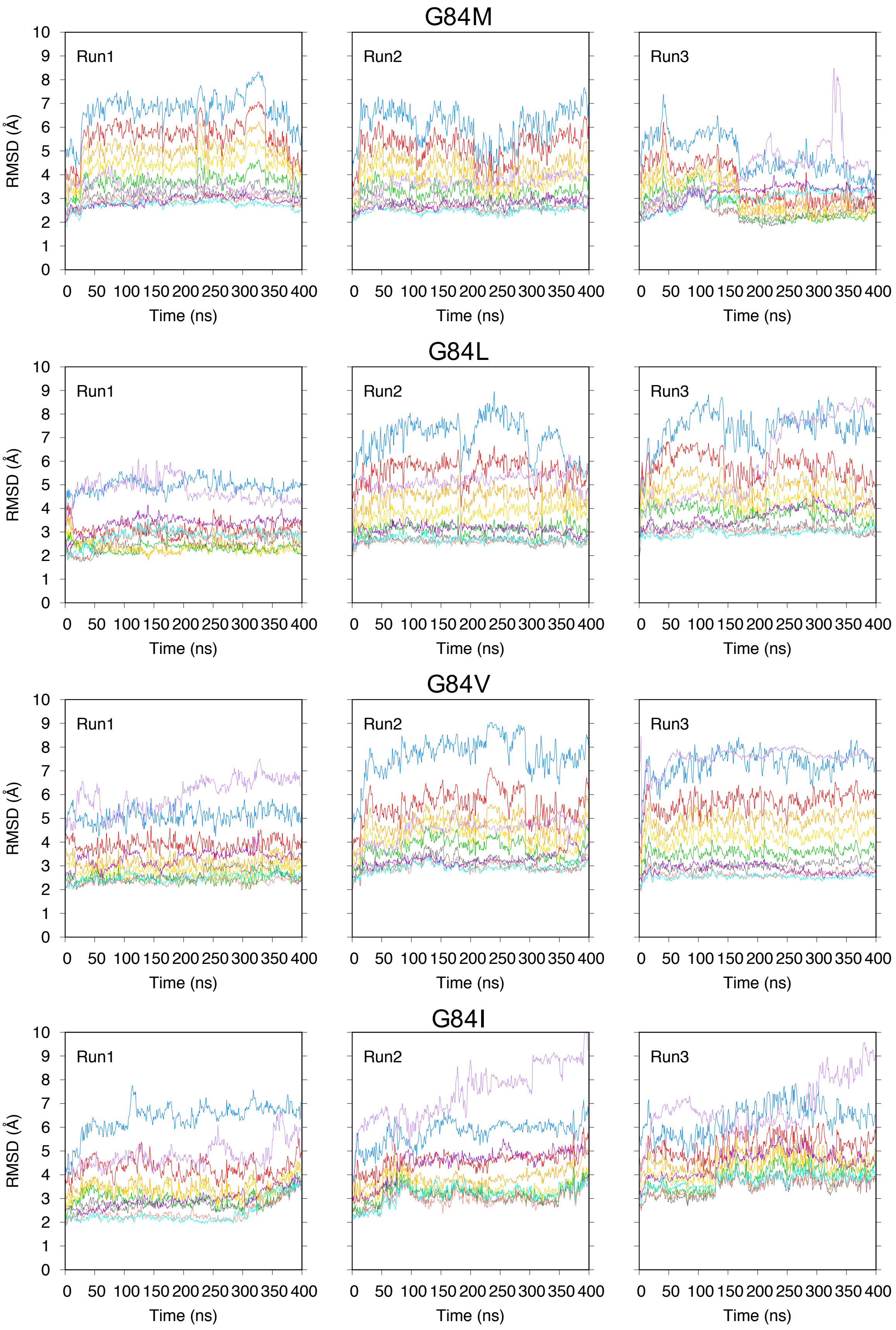


Figure S8: (cont'd.)

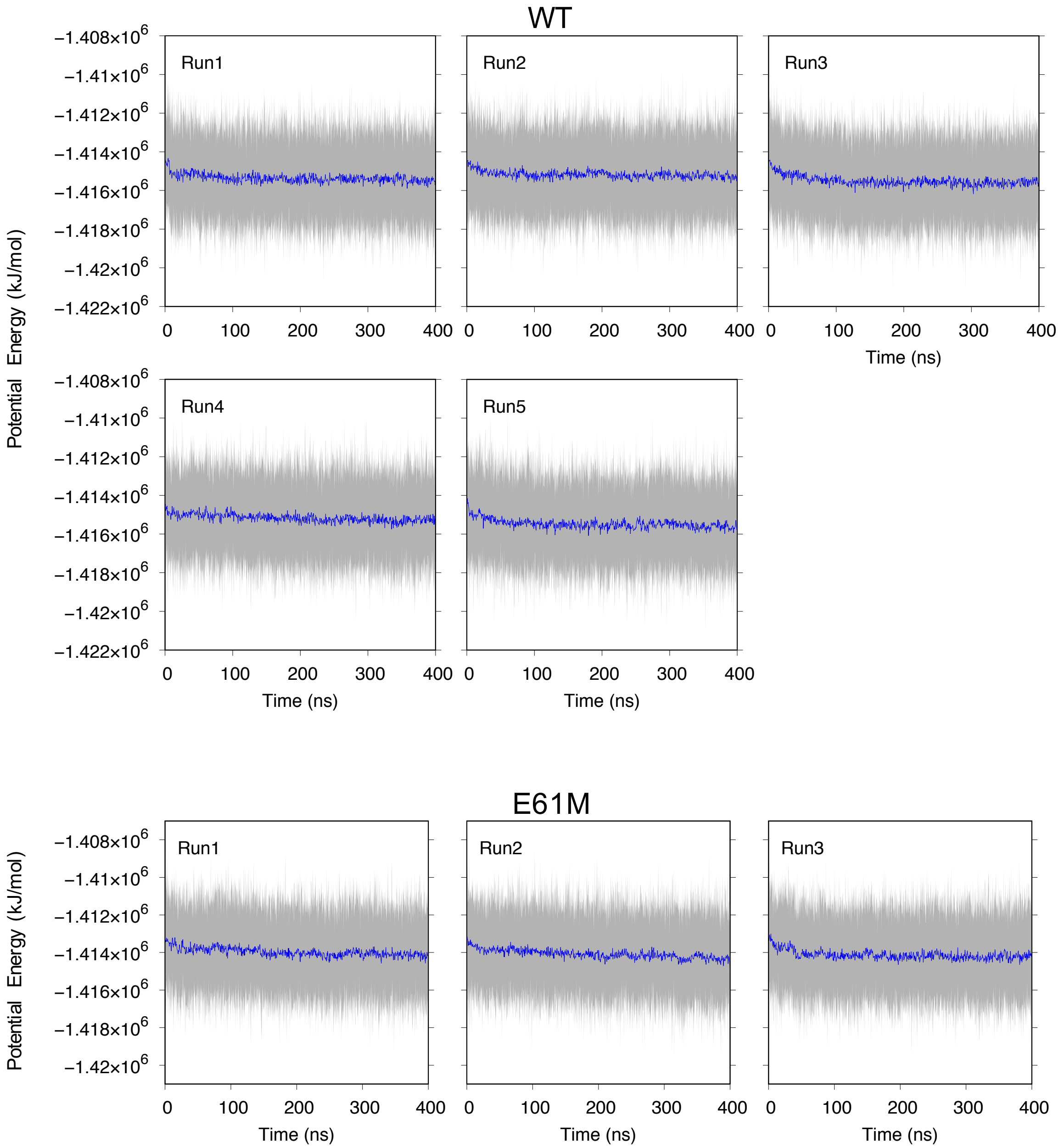


Figure S9: Potential energy as a function of time for the $\alpha$Syn amyloids (gray). The blue line represents a running average over 500 ps.

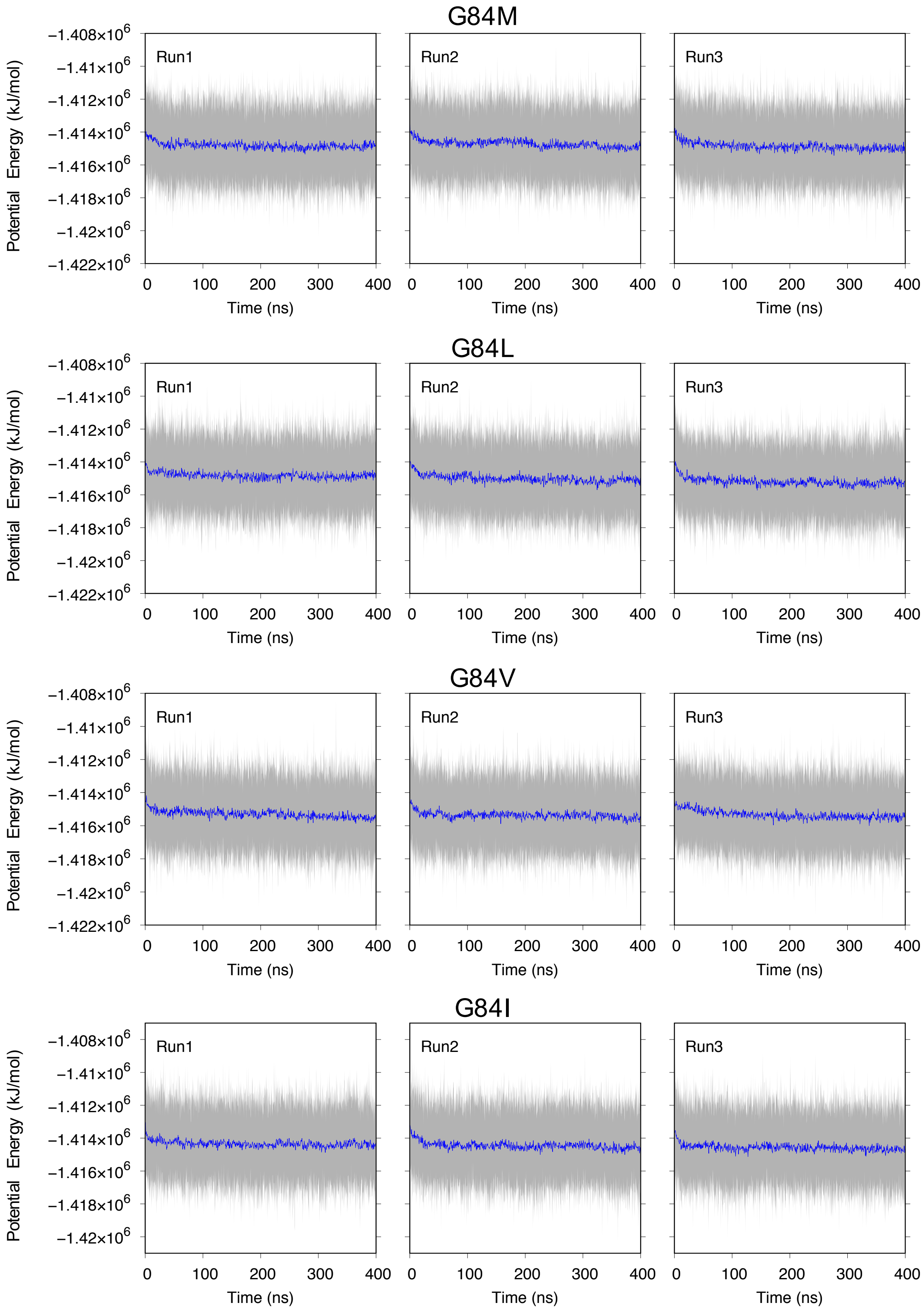


Figure S9: (cont'd.)

Table S2: Identical to Table S1, except for $PrP_{107\text{-}143}$

| Run | WT | | G127V | | M129V | |
|---|---|---|---|---|---|---|
| | RMSIP | Cumulative % | RMSIP | Cumulative % | RMSIP | Cumulative % |
| 1 | 0.69 | 79.9 | 0.67 | 71.3 | 0.69 | 75.0 |
| 2 | 0.61 | 75.8 | 0.70 | 66.3 | 0.65 | 75.0 |
| 3 | 0.70 | 79.8 | 0.65 | 73.5 | 0.65 | 67.6 |
| 4 | 0.68 | 75.2 | 0.66 | 78.0 | 0.71 | 74.1 |
| 5 | 0.71 | 77.2 | 0.68 | 68.2 | 0.65 | 73.8 |

| Run | I138M | | M129V&I138M | | A133V | |
|---|---|---|---|---|---|---|
| | RMSIP | Cumulative % | RMSIP | Cumulative % | RMSIP | Cumulative % |
| 1 | 0.70 | 75.4 | 0.64 | 66.5 | 0.67 | 76.9 |
| 2 | 0.70 | 85.6 | 0.66 | 74.7 | 0.68 | 75.8 |
| 3 | 0.68 | 79.8 | 0.66 | 79.0 | 0.75 | 70.7 |
| 4 | 0.70 | 69.8 | 0.62 | 75.6 | 0.72 | 76.3 |
| 5 | 0.66 | 71.2 | 0.73 | 63.1 | 0.64 | 84.0 |

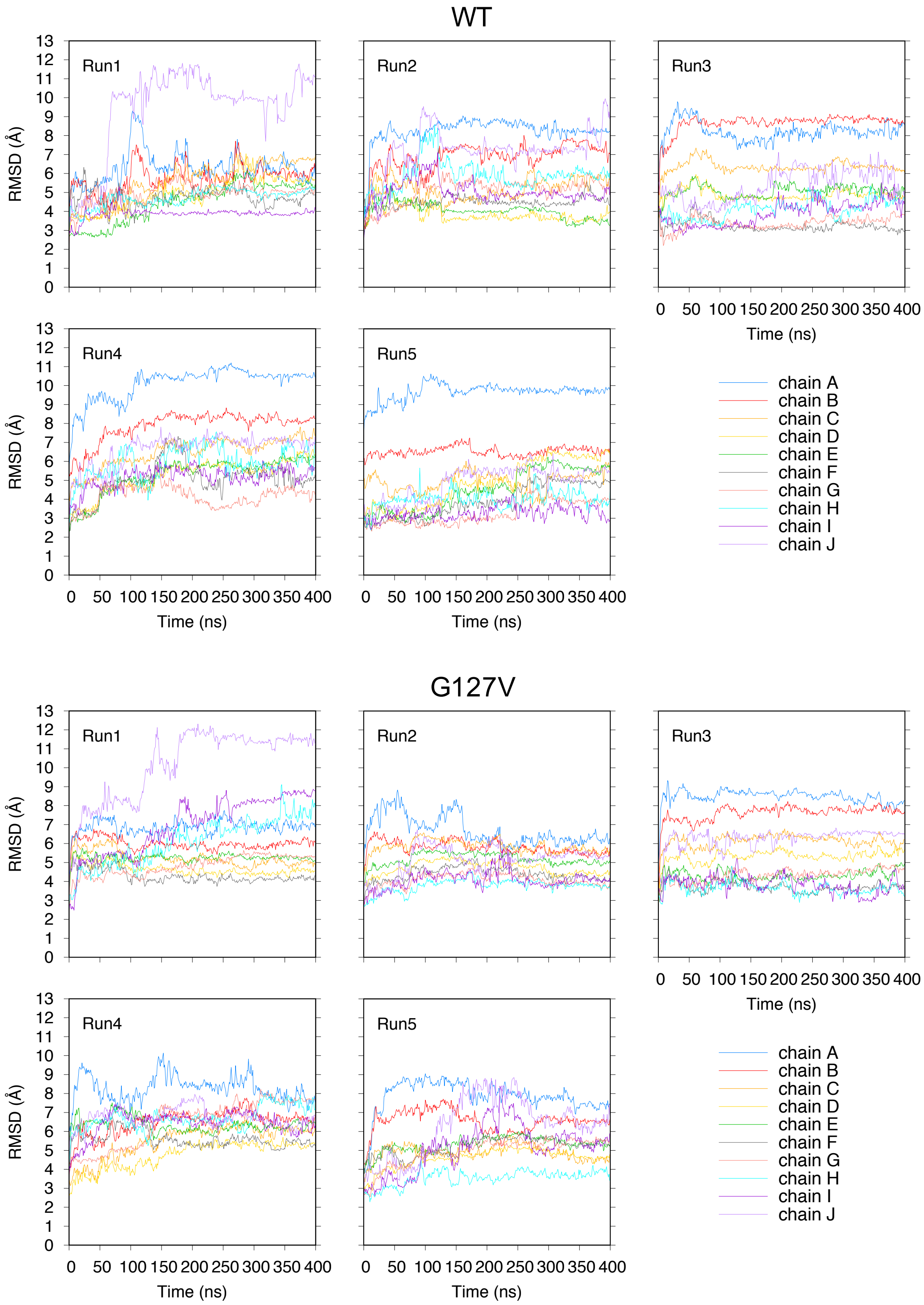


Figure S10: Backbone RMSD of $PrP_{107\text{-}143}$ protofibrils with respect to the constructed model (see Figure 4A).

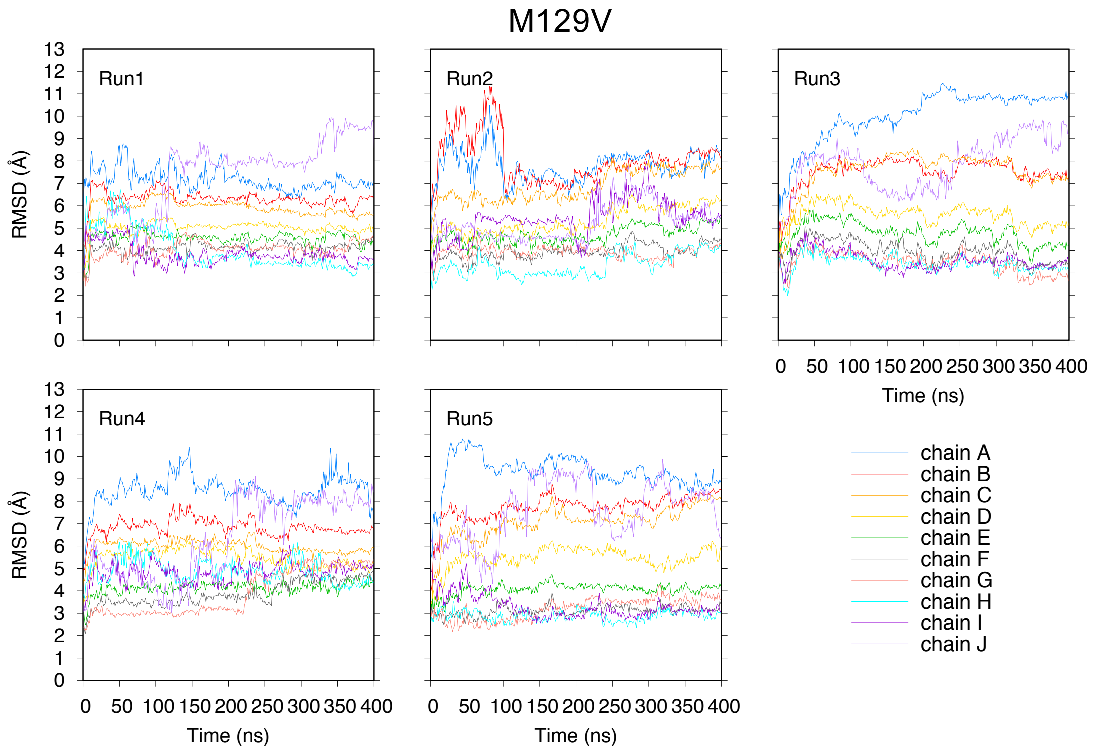


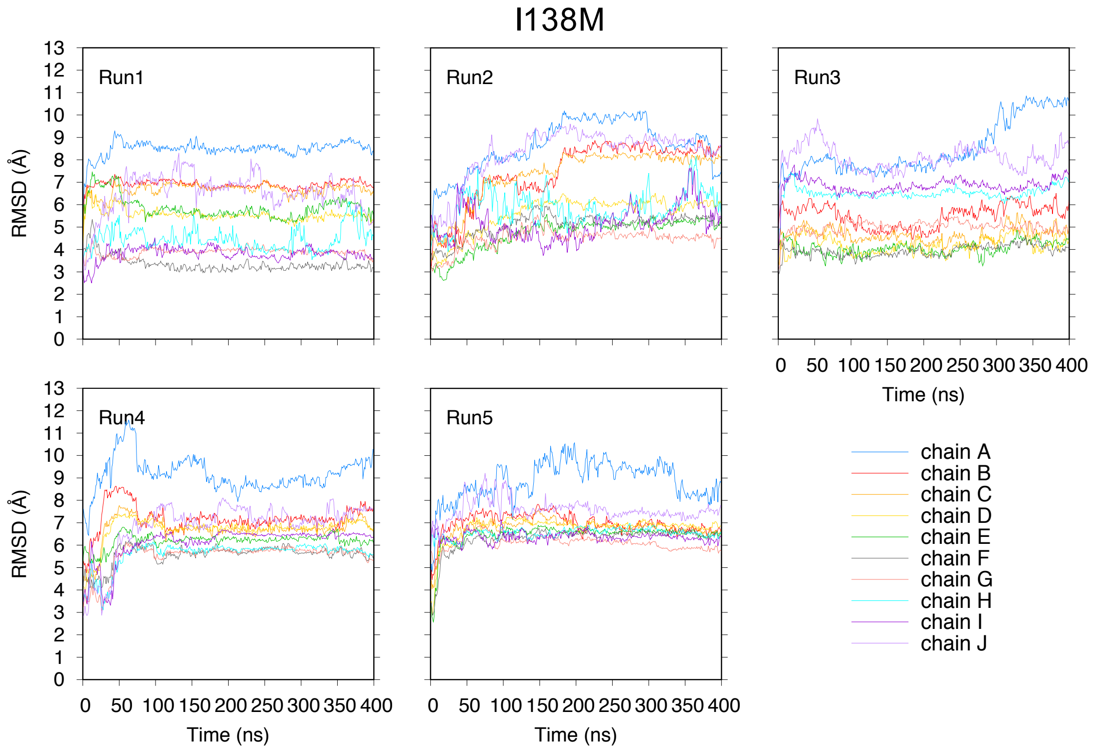


Figure S10: (cont'd.)

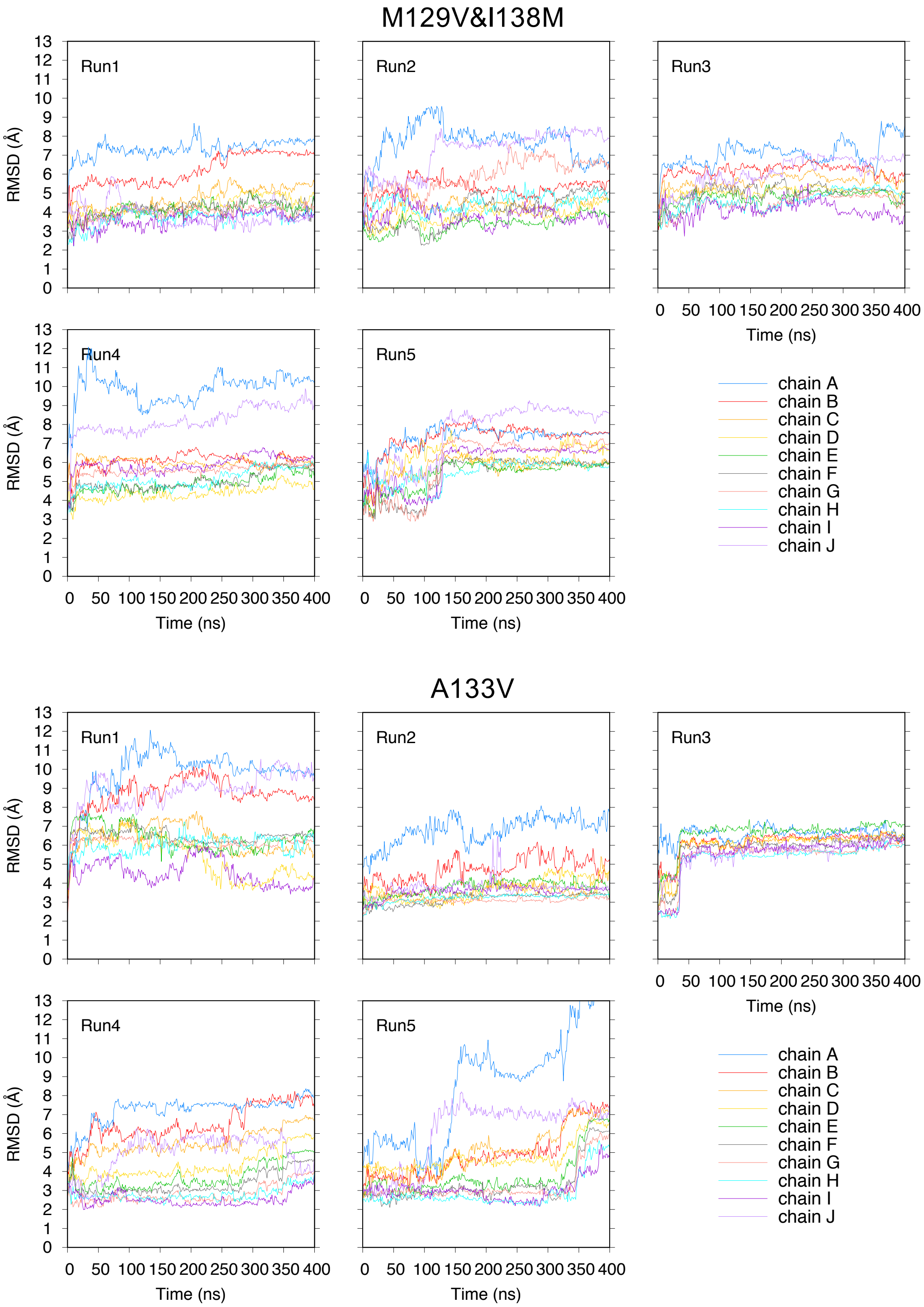


Figure S10: (cont'd.)

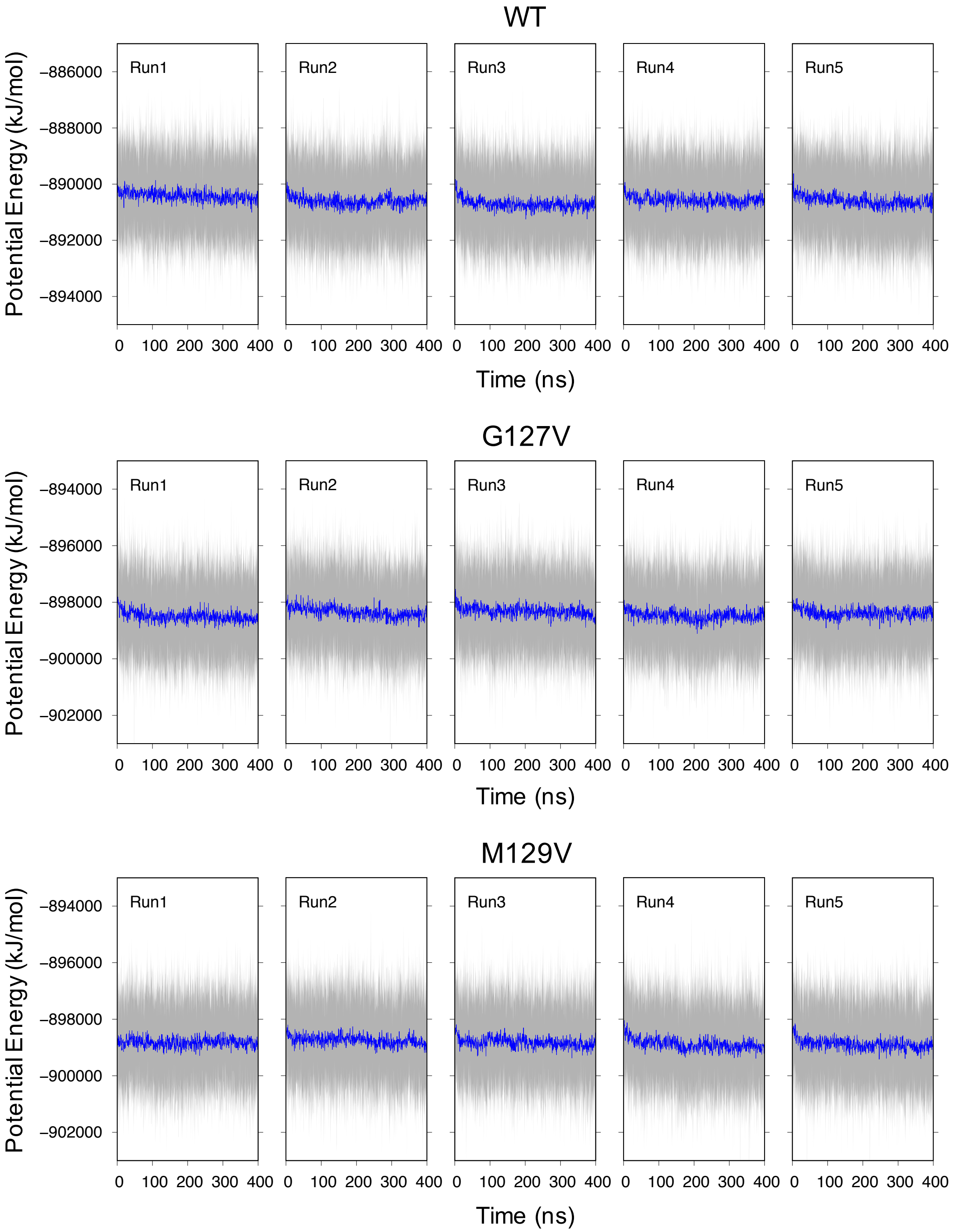


Figure S11: Potential energy as a function of time for $PrP_{107\text{-}143}$ protofibrils (gray). The blue line represents a running average over 500 ps.

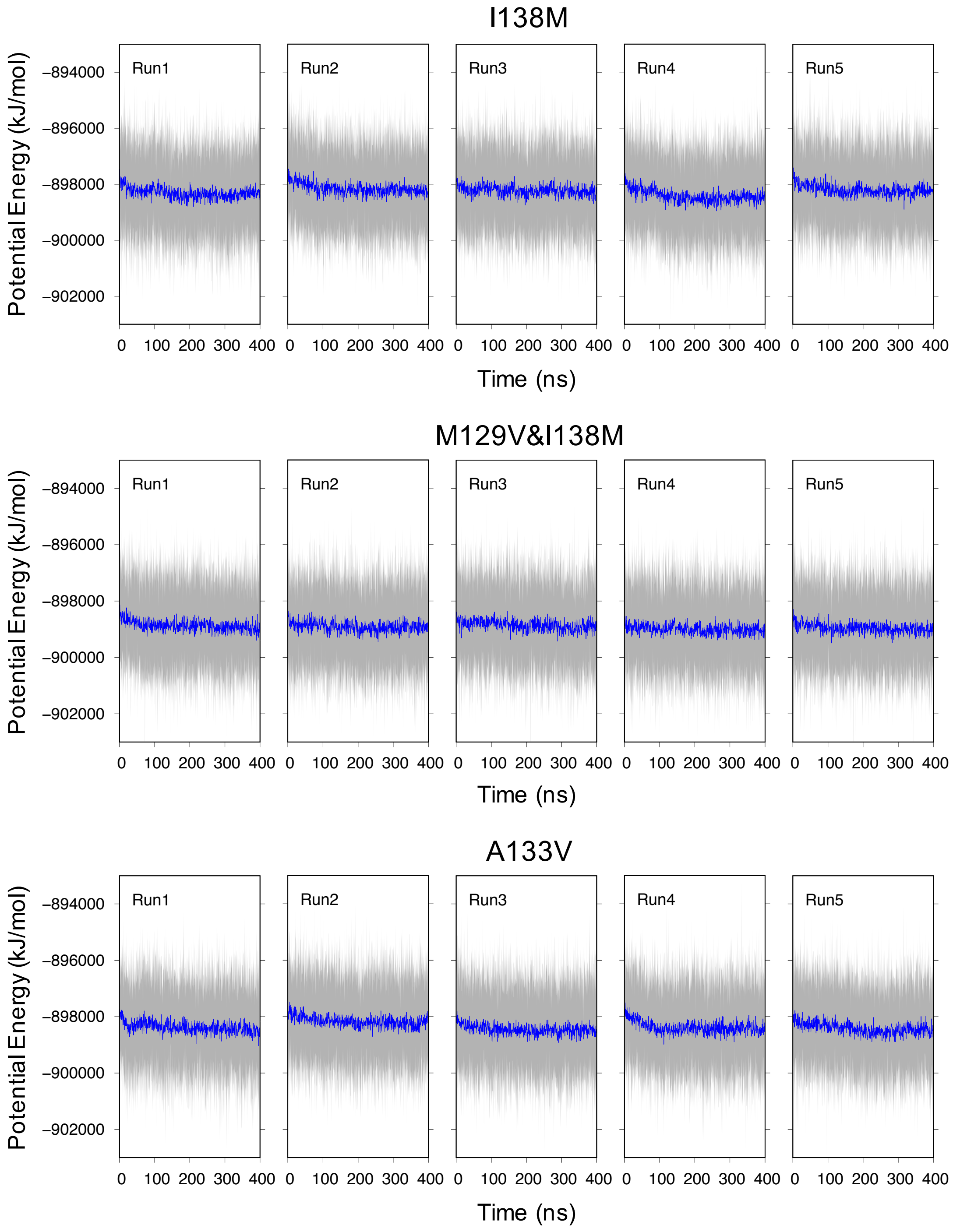


Figure S11: (cont'd.)
